# Light adaptation largely preserves the relative organization of parallel retinal channels: implications for efficient coding theory

**DOI:** 10.1101/2022.09.15.508164

**Authors:** Kiersten Ruda, Andra M. Rudzite, David St-Amand, Na Young Jun, John Pearson, Greg D. Field

## Abstract

Two major functions performed by the retina are to establish the parallel processing of visual information and to adapt visual encoding to the trillion-fold range of light intensities encountered in the environment. Previous work has highlighted many specialized cell types and circuits that instantiate parallel processing and light adaptation. However, fully understanding either process requires identifying how light adaptation and parallel processing interact. One possibility is that light adaptation causes uniform or proportional scaling to the receptive fields (RFs) of different retinal ganglion cell (RGC) types, the output neurons of the retina. Alternatively, light adaptation could cause a reorganization of RF structures across RGC types. To resolve these possibilities, we examined how the spatiotemporal RF structure of six simultaneously measured RGC types in the rat retina change from rod- to cone-mediated light levels. While light adaptation altered the RF properties of all six RGC types, we found that the relative structure across different RGC types was largely preserved across light levels. Surprisingly, most RGC types retained their center-surround RF structure even at low light levels, an observation that is at odds with prior efficient coding predictions. However, we show these predictions are incomplete and when RFs interact over a finite viewing area, efficient coding predicts the retention of surrounds under low signal-to-noise conditions.

## Introduction

Visual processing in the mammalian retina culminates into 20-40 retinal ganglion cell (RGC) types that convey distinct information about the visual environment to the brain (Baden et al. 2016; Dacey 2004; Sanes and Masland 2015; Bae et al. 2018; Field and Chichilnisky 2007; Goetz et al. 2022). In addition to establishing this parallel processing, the retina is largely responsible for adapting to the trillion-fold range of light levels encountered between night and day in terrestrial environments (Shapley and Enroth-Cugell 1984). This adaptation is critical to avoid saturation while maintaining sensitivity. However, light adaptation in the retina is not limited to changes in gain; experiments in many species have demonstrated changes in both the spatial and temporal receptive fields (RFs) of RGCs, which describe the visual features encoded by a cell (Barlow, Fitzhugh, and Kuffler 1957; Enroth-Cugell and Shapley 1973; Troy, Bohnsack, and Diller 1999; Tikidji-Hamburyan et al. 2015; Grimes, Schwartz, and Rieke 2014; Wienbar and Schwartz 2022). For example, previous studies have shown that temporal integration is longer under rod-mediated conditions and shorter under cone-mediated conditions (Jakiela and Enroth-Cugell 1976). In the spatial domain, the strength of the antagonistic surround generally increases from rod to cone-mediated light levels (Barlow, Fitzhugh, and Kuffler 1957; Atick and Redlich 1992; Srinivasan, Laughlin, and Dubs 1982). There are also reports that the contrast preference of RGCs can change with light adaptation (Tikidji-Hamburyan et al. 2015). Such observed changes have not been systematically tracked across distinct RGC types, raising the question of how RGC functional diversity depends on the mean light level. There are many possibilities for how light adaptation impacts different RGC types, and they set different constraints on adaptation mechanisms and on how downstream circuits need to process input from multiple RGC types.

Given previous studies, there are at least four possibilities for how functional diversity across RGC types could depend on light level (**Figure 1**). One possibility is that spatial and/or temporal integration changes proportionally across cell types (**Figure 1A**) (Barlow, Fitzhugh, and Kuffler 1957; Chan, Freeman, and Cleland 1992). For example, a 50% increase in RF area in one type is accompanied by a similar change in RF areas across the other types. We call this option ‘proportional adaptation,’ and it would result in a preserved functional organization across cell types despite changes in individual types. A second possibility is that spatial and/or temporal RF structure becomes more similar across different cell types at low light levels, for example, and more divergent at high light levels (**Figure 1B**) (Jakiela and Enroth-Cugell 1976; Devries and Baylor 1997). We call this ‘inhomogeneous adaptation,’ and it would result in either a contraction or expansion of functional diversity across light levels. More similar RGC responses may be advantageous at starlight conditions, where redundant signals could combat low signal-to-noise ratios. A third possibility is a reorganization of encoding that could include some cell types exhibiting minimal changes in RF structure while others exhibit large changes (**Figure 1C**) (Tikidji-Hamburyan et al. 2015; Cowan et al. 2017). We call this ‘reorganizing adaptation.’ While functional diversity could be maintained under this adaptation scheme, it would likely require complex mechanisms to accurately decode retinal signals at different ambient light levels. Finally, light adaptation across RGC types could include some combination of the above forms, which we call ‘mixed adaptation’ (**Figure 1D**).

**Figure 1:**
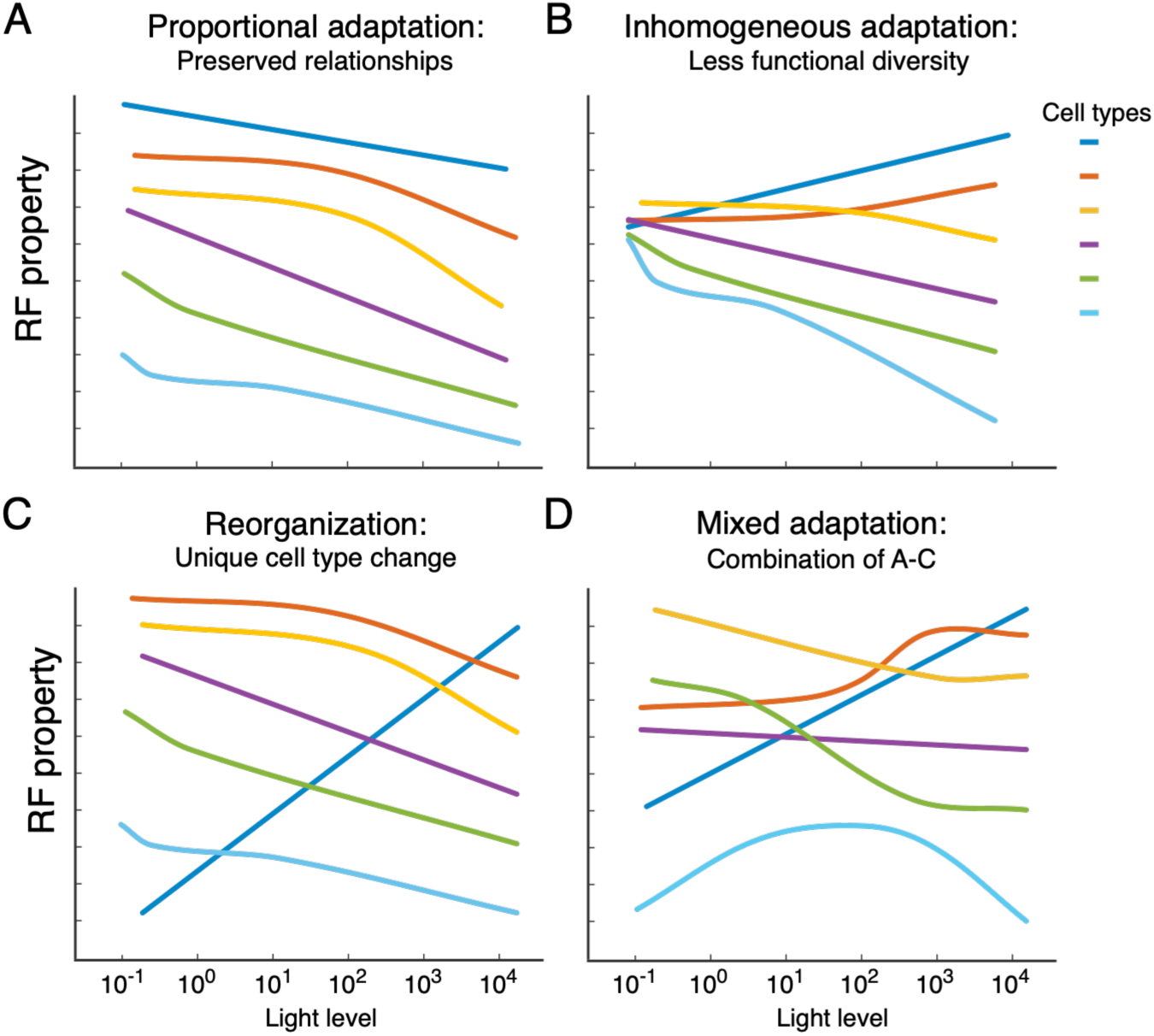
Possibilities for how adaptation affects the RFs and responses of different RGC types. (**A**) Proportional changes in different cell types would result in a stable relative organization of types across light levels. (**B**) Response properties of different types could become more similar at low light conditions. (**C**) Some cell types could exhibit little changes, while others show large changes. (**D**) Because the above options are not mutually exclusive, a fourth possibility is a combination of these adaptation effects.

To examine the impact of light adaptation on parallel processing, we performed large-scale multi-electrode array (MEA) measurements from ex vivo rat retina while presenting visual stimuli across light levels spanning rod- to cone-mediated vision. To make quantitative comparisons of RF properties across cell types, we focused on six RGC types exhibiting classical center-surround RFs that were nearly independent in space and time (Ravi et al. 2018). We measured the spatial and temporal RFs as well as the contrast response functions of these six types from starlight to sunlight conditions in 10-fold increments of light intensity. Our results show several invariances to the relative organization across the six RGC types. The relationships in spatiotemporal RFs and contrast responses across different types were largely preserved over light intensities, except for temporal or spatial integration in one cell type. These findings indicate ‘mixed adaptation’ across these RGC types that is composed mainly of ‘proportional adaptation.’ The dominance of proportional adaptation suggests that signals from RGC types can be stably processed by downstream circuits across rod- and cone-mediated conditions.

## Results

To understand how light adaptation alters RGC encoding at the level of populations rather than single cells, we consider changes in the relative visual features encoded by different RGC types. We measured the spiking activity of hundreds of rat RGCs from rod-mediated (scotopic) to cone-mediated (photopic) conditions using large-scale multi-electrode arrays (MEAs) (Litke et al. 2004; Frechette et al. 2005; Ravi et al. 2018). Retinal samples were presented with visual stimuli whose mean light intensity spanned a range of 6 log units, from 0.1 to 10,000 Rh*/rod/s. Note that after switching light levels, retinas were allowed to adapt for at least 10 minutes. Thus, our recordings and analysis reflect the steady state of RGC responses at each light level rather than rapid adaptation (<60s) (Mahroo and Lamb 2004; Cameron et al. 2008; Joachimsthaler, Tsai, and Kremers 2017; Wark, Fairhall, and Rieke 2009).

We functionally classified RGCs recorded at 10,000 Rh*/rod/s using response properties such as direction selectivity, the shape of the spike train autocorrelation, and receptive field (RF) characteristics like temporal integration and spatial RF size (Ravi et al. 2018). The accuracy of cell typing was confirmed by a spatial arrangement of RFs within each type that forms a mosaic-like tiling of space (**Figure 2**) (Devries and Baylor 1997; Field et al. 2007; Ravi et al. 2018). We focus on six RGC types whose RFs have a classical center-surround organization (Kuffler 1953) (**Figure 2**). The types are named based on differences in their response polarities (ON or OFF) and dynamics of temporal integration (Brisk Sustained, Brisk Transient, or Transient). While not definitive, these RGC types have likely homologies in other species. ON-brisk sustained (ON-bs) RGCs may be homologous to ON-sustained alpha RGCs (a.k.a ON-delta) in other rodents and ON midget RGCs in primates (likewise for OFF-bs RGCs) (Peichl 1989). Similarly, ON-brisk transient (ON-bt) RGCs are likely homologous to ON-transient alpha RGCs in other rodents and ON parasol RGCs in primates (Ravi et al. 2018; Crook et al. 2008; Krieger et al. 2017; Hahn et al. 2023). Below, we systematically analyze changes to the features encoded by each cell type, beginning with RF structure.

**Figure 2:**
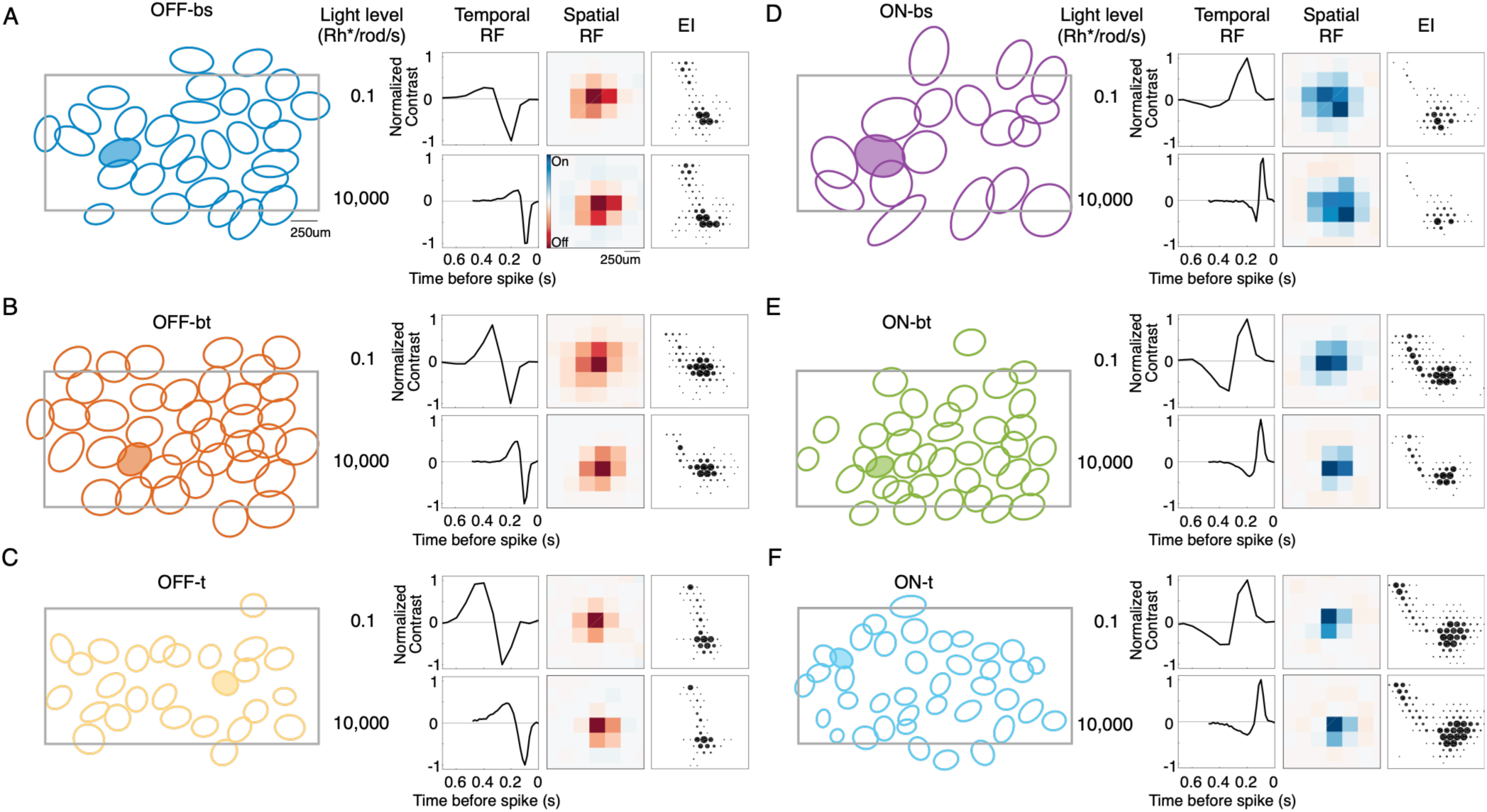
Cell types and representative RFs across light levels. (**A-F**) Left: Mosaics formed by the RFs of each RGC in a type. Right: Temporal RF, spatial RF, and electrophysiological image (EI) of highlighted cell from Left at a scotopic (top) and photopic (bottom) light level. See also Figure 2-supplemental figure 1. Data from one retina.

RFs were estimated by presenting checkerboard noise and calculating the spike-triggered average (STA) at each light level (**Figure 2**) (Chichilnisky 2001). As RF properties changed across light levels, they could not be used to track individual cells reliably. We used two strategies to ensure RFs of a cell at one light level were compared to the same cell at other light levels: 1) principal components analysis of spike waveform shapes (see Methods; (Yu et al. 2017)) and 2) the electrophysiological image (EI; see **Figure 2**) (Litke et al. 2004). The EI is generated by computing the spike-triggered mean electrical activity over the MEA, which generates an electrical ‘footprint’ of each RGC (Petrusca et al. 2007). This information allows for the reliable tracking of individual RGCs across light levels spanning rod to cone vision (Field et al. 2009; Yao et al. 2018). In what follows, we separately analyzed the temporal and spatial RFs of each cell type across light levels. The spatial and temporal RFs summarize the linear component of spatial and temporal integration, respectively. All six cell types analyzed exhibited spatiotemporal RFs that were closely approximated by the outer product of a single spatial RF and single temporal RF, which justified separately analyzing each component (**Figure 3** and **Figure 3 - supplemental figure 1**, see Methods) (DeAngelis, Ohzawa, and Freeman 1993).

**Figure 3:**
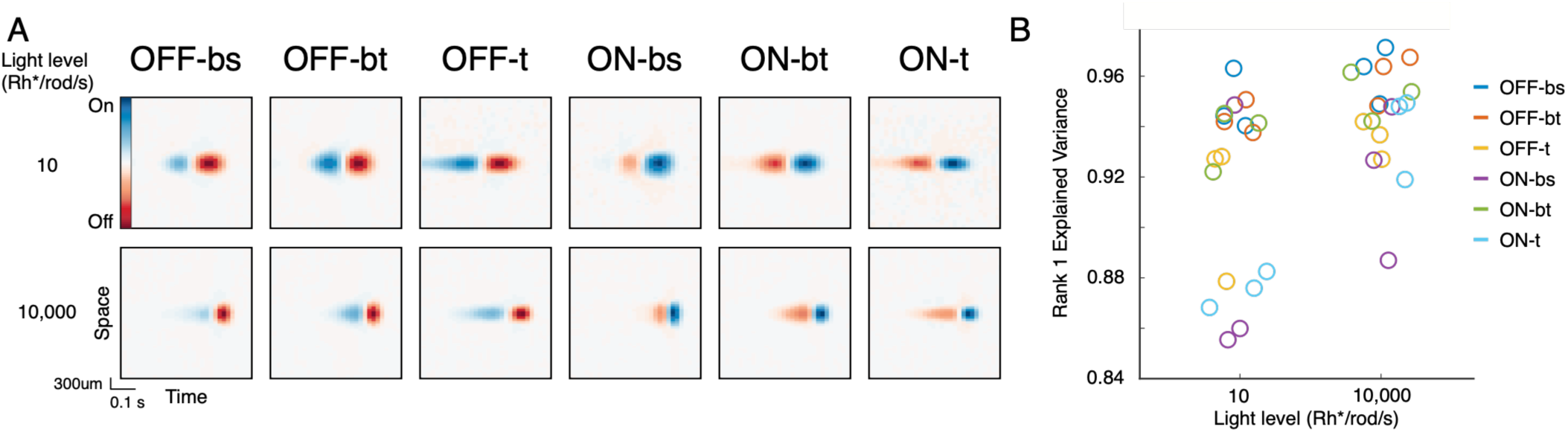
RFs are largely space-time separable. (**A**) Space-time view of average RFs from one retina at two light levels. The plots show a cross-section of the spatial RF at time preceding the spike (right edge of each plot). The weak diagonal structure indicates little interaction between space and time. (**B**) RF variance explained by a single pair of spatial and temporal filters (rank 1). Each point is the explained variance in the average STA for a given cell type and light level in one retina (3 retinas total). The total number of RGCs depended on light level. For OFF-bs, there were 75 and 95 recorded cells at the scotopic and photopic light level, respectively; OFF-bt: 122 and 131; OFF-t: 18 and 27; ON-bs: 51 and 66; ON-bt: 74 and 84; ON-t: 14 and 29. See also Figure 3-supplemental figure 1.

### The relative structure of temporal integration is maintained, except for ON-bs RGCs

We first examined changes in temporal RFs across RGC types (**Figure 4**). All six types exhibited a substantial temporal compression with increasing light levels (**Figure 4A**). For example, OFF-t RGCs integrated visual input over a ∼600 ms time window at 0.1 Rh*/rod/s, which decreased to less than 400 ms at 10,000 Rh*/rod/s (p<0.001 for all types). While this compression is expected in the switch from rod- to cone-mediated vision (Enroth-Cugell and Shapley 1973; Purpura, Kaplan, and Shapley 1988), we were primarily interested in whether the relative changes across cell types exhibited proportional, inhomogeneous, reorganizing, or mixed adaptation (**Figure 1**). Comparing the duration of temporal integration across types shows that the relative relationships are approximately maintained (**Figure 4B**). OFF-t and ON-t have some of the slowest integration at every light level, the OFF-bt type exhibits fast integration, and ON-bt is in the middle (ON-bt zero crossing is smaller than OFF-t at all light levels except 0.1 Rh*/rod/s, p<0.05; ON-bt zero crossing is larger than OFF-bt at all light levels, p<0.001). Thus, the duration of temporal integration is largely preserved from rod- to cone-mediated conditions. A major exception to this trend is the adaptation among ON-bs RGCs (**Figure 4B**, purple). At 10,000 Rh*/rod/s, this cell type exhibits the briefest integration (faster than all other types, p<0.001). However, the ON-bs RGCs slow proportionally more than the other types at lower light levels (**Figure 4C**, p<0.001 comparing percent change from 0.1 to 10,000 Rh*/rod/s). Thus, the duration of temporal integration exhibits mixed adaptation that is dominated by proportional adaptation across five of six RGC types. The absolute area under the temporal RFs indicates a similar trend: the ON-bs RGCs exhibit some of the largest fractional changes in area (**Figure 4E**, p<0.05 compared to all types except ON-t and OFF-t).

**Figure 4:**
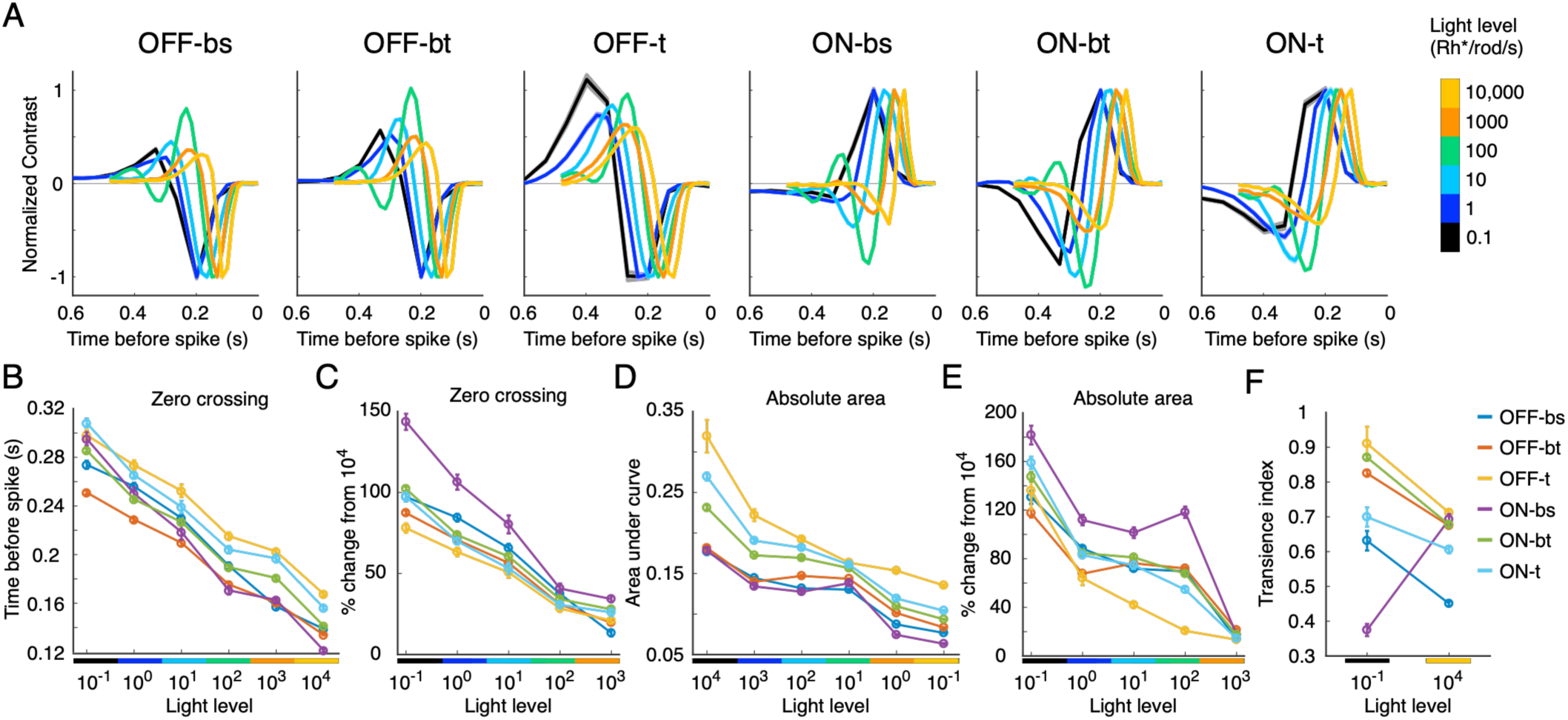
Relationships among temporal RFs of different types are relatively stable over light levels, with the exception of ON-bs RGCs. (**A**) Average temporal RFs from one retina organized by cell type (left to right). Thickness of curves indicates SEM. (**B**) Zero cross in temporal RFs across light levels, as measured by the time the temporal RF crosses the x-axis. (**C**) Percent change of zero crossing. ON-bs time courses slow proportionally more at scotopic conditions. (**D**) Absolute area under temporal RFs. The relative ordering of absolute area is maintained over light intensities. (**E**) Percent change of temporal RF area. Error bars are SEM. (**F**) Degree of transience for the lowest and highest light levels. All panels except for A accumulate data from four retinas. The total number of RGCs depended on light level. For OFF-bs, the range of recorded RGCs over all light levels is 34-107; OFF-bt: 48-160; OFF-t 10-63; ON-bs 20-51; ON-bt 52-130; ON-t 16-64. See also Figure 4-supplemental figure 1.

We next assessed how light adaptation alters the dynamics or shape of temporal RFs, which determines the stimulus kinetics encoded by the RGC. For instance, a temporal RF with a single lobe (monophasic) will produce a sustained response to a step of light. This integration is equivalent to a low pass filter in the frequency domain (see **Figure 5**). Conversely, a temporal RF with a positive and negative lobe (biphasic) will produce a transient response to a step of light, signaling a temporal change. This is equivalent to a band pass filter in the frequency domain (Purpura, Kaplan, and Shapley 1988). Theoretical studies predict temporal tuning changes from biphasic at high light levels to monophasic at low light levels (Atick and Redlich 1990, 1992; van Hateren 1992; Srinivasan, Laughlin, and Dubs 1982). The theory suggests that RGC computations are limited by stimulus noise under scotopic conditions, shifting from computing a temporal difference at high light levels to pure integration at low light levels. To test this prediction across many cell types, we computed a transience index that ranges from 0 (sustained) to 1 (biphasic, with an equal area under both lobes; see Methods). Only ON-bs RGCs followed the theoretical prediction, becoming much less transient as the light level decreased (**Figure 4F**, purple; p<0.001 comparing 10,000 and 0.1 Rh*/rod/s). Surprisingly, all other types exhibit the opposite trend, becoming more biphasic at scotopic light levels (p<0.001). For these five types, the relative ordering of transience indices was also stable (OFF-bt, OFF-t, and ON-bt are more transient than OFF-bs and ON-t at 0.1 and 10,000 Rh*/rod/s, p<0.001). Thus, mixed adaptation governs the shape of the temporal RFs, but five of the six types exhibit approximately proportional adaptation.

**Figure 5:**
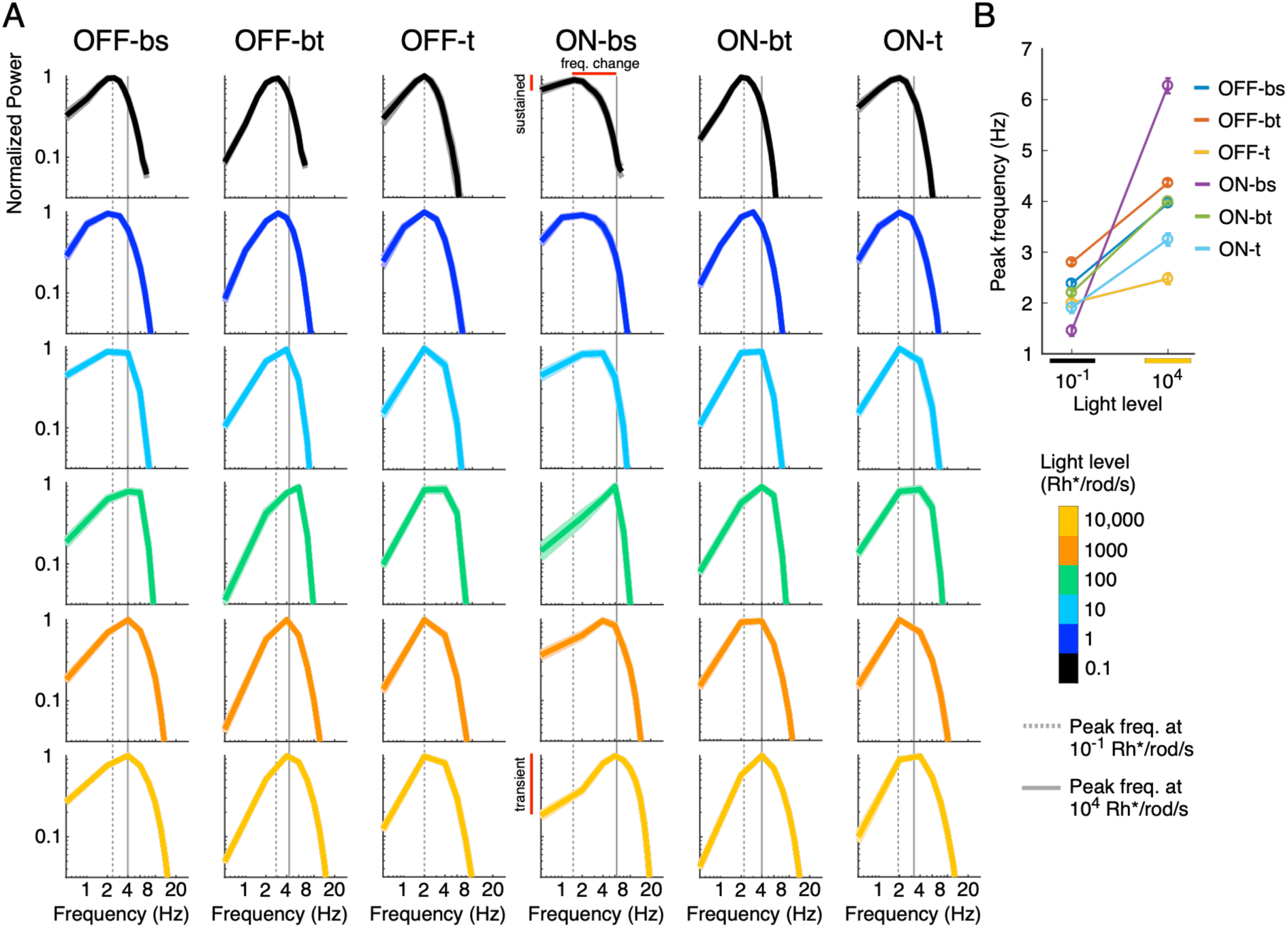
Temporal frequency tuning changes across light levels. (**A**) Normalized temporal frequency tuning averaged over four retinas for each cell type and light level. Note that the ON-bs curves become low pass at 0.1 Rh*/rod/s, while all other types remain relatively band pass (see also Fig 4F). Line thickness indicates SEM. (**B**) Peak frequency of temporal RFs demonstrates that ON-bs RGCs exhibit slowed integration compared to the other cell types. The total number of RGCs is the same as in Figure 4.

To inspect the filtering properties of temporal RFs more explicitly, we replot them in frequency space (**Figure 5A**) (Shapley and Enroth-Cugell 1984). This view illustrates that ON-bs tuning becomes more low-pass with decreasing light levels, while all other types remain band-pass. We also extracted the peak temporal frequency of these curves to assess changes in the frequency domain. As light intensity decreases, the peak frequency decreases for all six RGC types (**Figure 5B**), reflecting the prolonged temporal integration found in **Figure 4** (peak frequency at 10,000 Rh*/rod/s is larger than that at 0.1 Rh*/rod/s, p<0.001 for all types except OFF-t where p=0.1). Again, ON-bs RGC tuning slows proportionally more than the other types, going from the fastest peak temporal frequency at 10,000 Rh*/rod/s (p<0.001) to one of the slowest at 0.1 Rh*/rod/s (slower than ON-bt, OFF-bs, and OFF-bt, p<0.001) with a large percent change in peak frequency (77 +/- 2% decrease from 10,000 Rh*/rod/s).

These temporal RF changes show several departures from previous work. First, we find that most cell types remain band-pass under scotopic conditions, indicating that stimulus noise is not a limiting factor for detecting temporal contrast at those light levels (Atick and Redlich 1990, 1992; van Hateren 1992; Brinkman et al. 2016). Second, the relative ordering of temporal RFs is mostly maintained across cell types except for ON-bs RGCs, which exhibit distinct changes to their temporal RF duration and dynamics (see Discussion).

### Spatial RFs increase in size but retain surrounds at scotopic conditions

We next turn to analyze the dependence of spatial RFs on light level. Many previous studies have estimated changes in spatial RFs across light levels, with varied results (Barlow, Fitzhugh, and Kuffler 1957; Troy, Oh, and Enroth-Cugell 1993; Troy, Bohnsack, and Diller 1999; Field et al. 2009; Cowan et al. 2017). The canonical view is that the antagonistic surround becomes weaker while the RF center expands at low light levels (Barlow, Fitzhugh, and Kuffler 1957). This change corresponds to a switch from band-pass spatial filtering in photopic conditions to low-pass spatial filtering in scotopic conditions, consistent with theoretical predictions described above (Atick and Redlich 1992).

We first examined changes in RF center size across light levels using checkerboard noise stimuli (**Figure 6**). These measurements were limited to light levels of 1.0 Rh*/rod/s and above because relatively small checkers were required to accurately estimate the RF center size and measure small changes in RF center size across light levels. At lower light levels (i.e., ∼0.1 Rh*/rod/s), Poisson variability in the number of photons absorbed is large relative to the contrast of the flickering checkers when the checkers are small (e.g., 120 µm). RF center size was estimated from the spatial component of the STA (**Figure 2**) by fitting a two-dimensional difference-of-Gaussians to each spatial RF. One Gaussian captured integration by the center, while the other Gaussian accounted for the RF surround. The center radius was slightly smaller under cone-mediated conditions for most cell types (**Figure 6B**, p<0.05 for OFF-bt, OFF-t, and ON-bt). The relative sizes were also mostly preserved across light levels (e.g., ON-t and OFF-t had the smallest RFs and ON-bs and OFF-bt had the largest RFs; OFF-t and ON-t centers are smaller than those of OFF-bt and ON-bs at each light level, p<0.005). The exception to proportional adaptation was inhomogeneous adaptation between OFF-bs and ON-bt RGC types, which exhibited similar RF sizes at the highest light level, but different sizes at the lowest light level (**Figure 6B**, green and blue, p<0.001).

**Figure 6:**
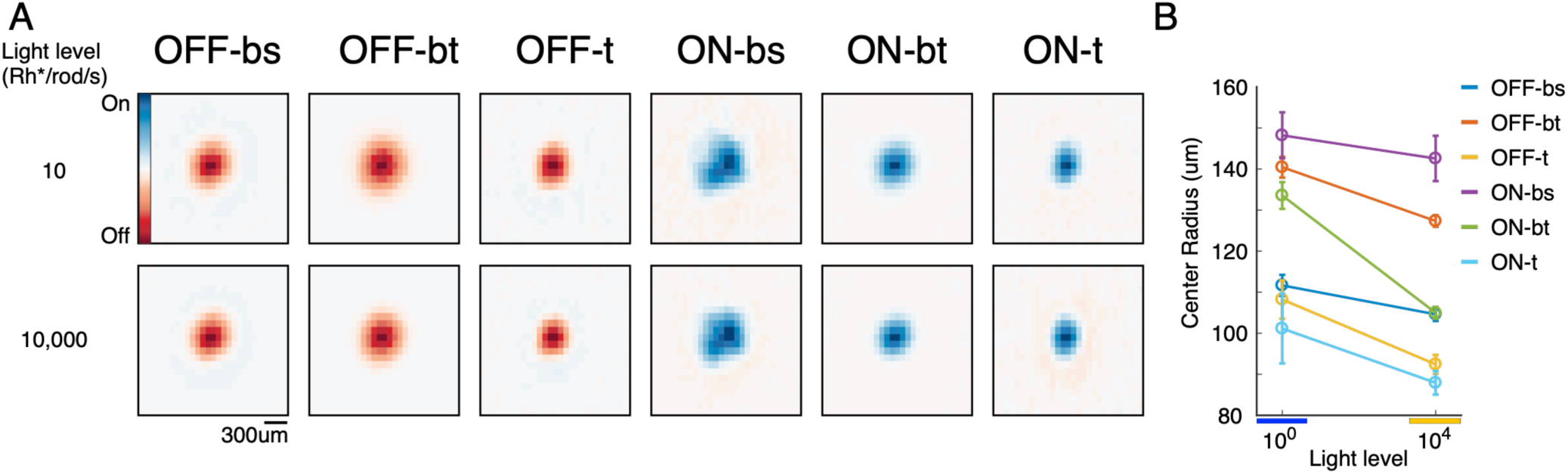
The center size of spatial RFs decreases with increasing light intensity. (**A**) Average spatial RF over all RGCs in one retina, shown at two light levels. (**B**) Comparison of average RF center size between scotopic and photopic conditions. RFs were measured with 63x63 µm stimulus pixels (checkers), and error bars show SEM. Data accumulated from three retinas. The total number of RGCs depended on light level. For OFF-bs, the range of recorded RGCs over all light levels is 39-97; OFF-bt: 52-132; OFF-t: 5-29; ON-bs: 5-66; ON- bt: 36-88; ON-t: 5-32.

A drawback to checkerboard noise is that it may not lead to accurate measurements of RF surrounds. The small checkers necessary to estimate center size can produce too low an effective contrast for estimating the much larger RF surround (Wienbar and Schwartz 2018). We therefore complemented our STA analysis of spatial RFs with sinusoidal drifting gratings that can more effectively drive RF mechanisms at low spatial frequencies (**Figure 7**). For each grating size, we extracted the response amplitude at the characteristic temporal frequency of the stimulus (2Hz). This generated a spatial frequency tuning function for each RGC across light levels and cell types (**Figure 7A**). Individual cell tuning functions were fit with a difference of Gaussians model to estimate the RF center size, surround size, and relative surround/center strength (Enroth-Cugell and Robson 1966). Consistent with the results from checkerboard noise (**Figure 6**), RF centers of each type expanded at the lowest light levels (**Figure 7B**, p<0.005). The prediction for RF surrounds from previous experiments (Barlow, Fitzhugh, and Kuffler 1957), and theoretical studies (Atick and Redlich 1992) indicate that surround strength decreases in scotopic conditions (but see (Troy, Bohnsack, and Diller 1999; Cowan et al. 2017)). In contrast, we observed robust RF surrounds at the lowest light levels we tested (0.1 Rh*/rod/s; **Figure 7C**). ON-bt RGCs were an exception; their RF surrounds were nearly absent at the lowest light level. Thus, these data depart from theoretical predictions and previous studies by demonstrating that strong surrounds are maintained across light levels in most (five of six) rodent RGC types with a classical center- surround RF structure.

**Figure 7:**
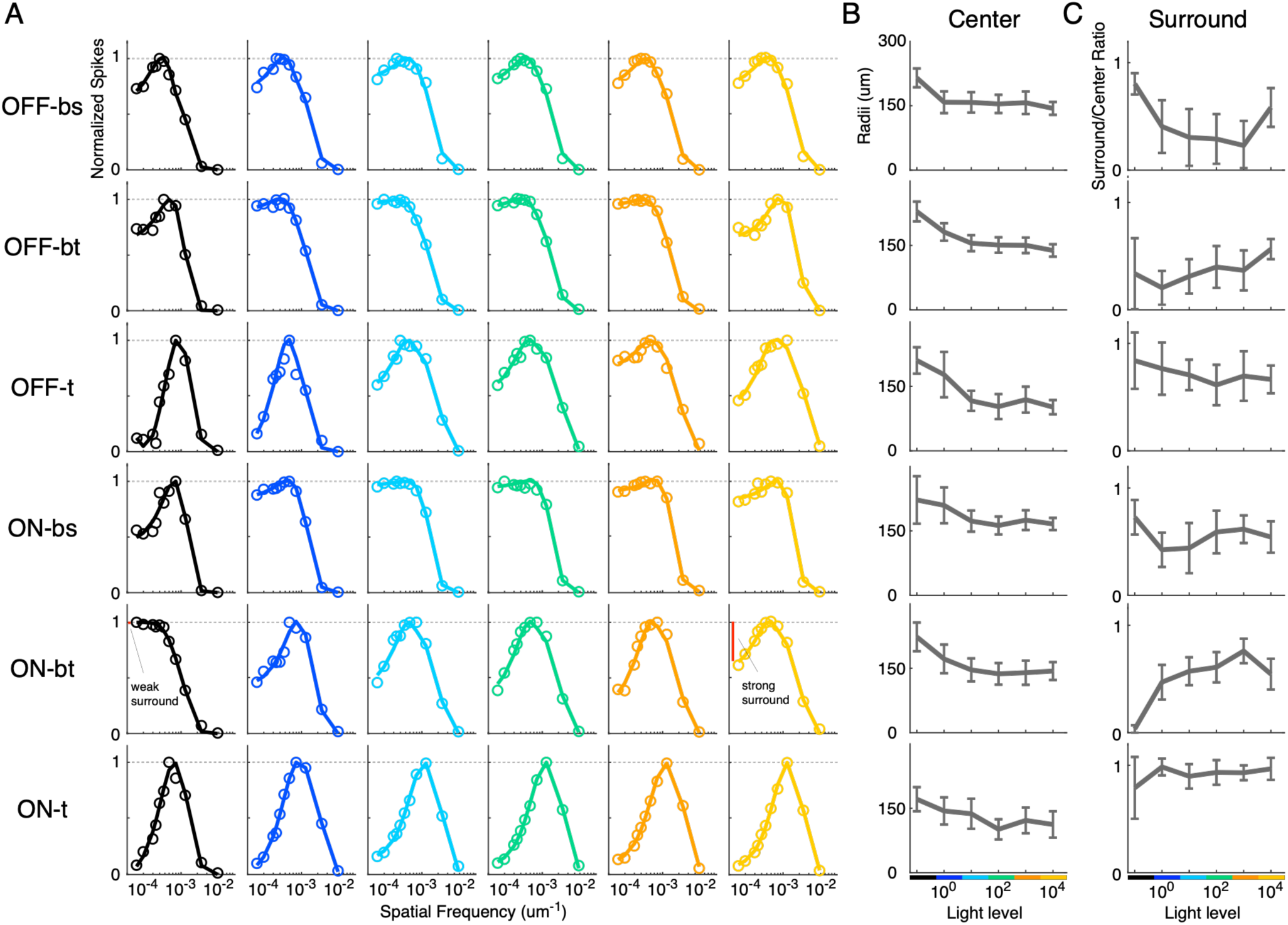
For most RGC types, RF surrounds remain robust in scotopic conditions. (**A**) Example spatial tuning curves across light levels for each cell type. The band pass shapes indicate that antagonistic surrounds are present in the spatial RFs. (**B**) Median and median absolute deviation (error bars) and S.E.M of RF center size across light levels for the 6 RGC types. (**C**) Median and median absolute deviation of surround-center ratios across light levels for the 6 RGC types. The total number of RGCs depended on light level. For OFF-bs, the range of recorded RGCs over all light levels is 31-42; OFF-bt: 52-72; OFF-t: 18-27; ON-bs: 26-37; ON-bt: 50-69; ON-t: 13-23.

### Spatial and temporal integration are inversely related at all light levels

Among RGC types of several species, including primates and rodents, temporal and spatial resolution trade-off for one another: cells with the largest spatial RFs exhibit the briefest temporal integration and vice versa (Frishman et al. 1987; Ravi et al. 2018; Ocko et al. 2018; Jun, Field, and Pearson 2022). Thus, a tradeoff between spatial and temporal resolution appears to be an organizing principle in the retina and perhaps elsewhere in the visual system (DeAngelis, Ohzawa, and Freeman 1993; De Valois et al. 2000). To determine how this space-time organization might change with light adaptation, we compared the spatial and temporal integration for the six cell types across light levels (**Figure 8**). A space-time tradeoff was present at all light levels, with similar slopes defining the tradeoff across light levels and cell types. Considering each RGC type individually (**Figure 8B**), resolution in both space and time decreases with light level, although not in a perfectly linear manner. To determine if the extent of tradeoff is stable over light levels, we aligned data across light levels with ordinate and abscissa offsets and examined how well a single line described the data (**Figure 8A**, red lines). The average R^2^ performance of the fixed slope is 0.51 (**Figure 8C**), indicating that the extent of the space-time tradeoff is approximately constant across light levels. One limiting factor for a single fixed slope to explain the data may be due to strong changes in ON-bs temporal integration: ON-bs temporal RFs slow substantially more than other cell types at lower light levels (**Figs. 4 & 5**). However, their spatial RF sizes are more constant across light levels (**Figs. 6 & 7**), which would not be expected under a constant space-time tradeoff. In summary, the space-time tradeoff across cell types and light levels is approximately consistent with proportional adaptation, with the ON-bs RGC likely contributing some reorganizing adaptation.

**Figure 8:**
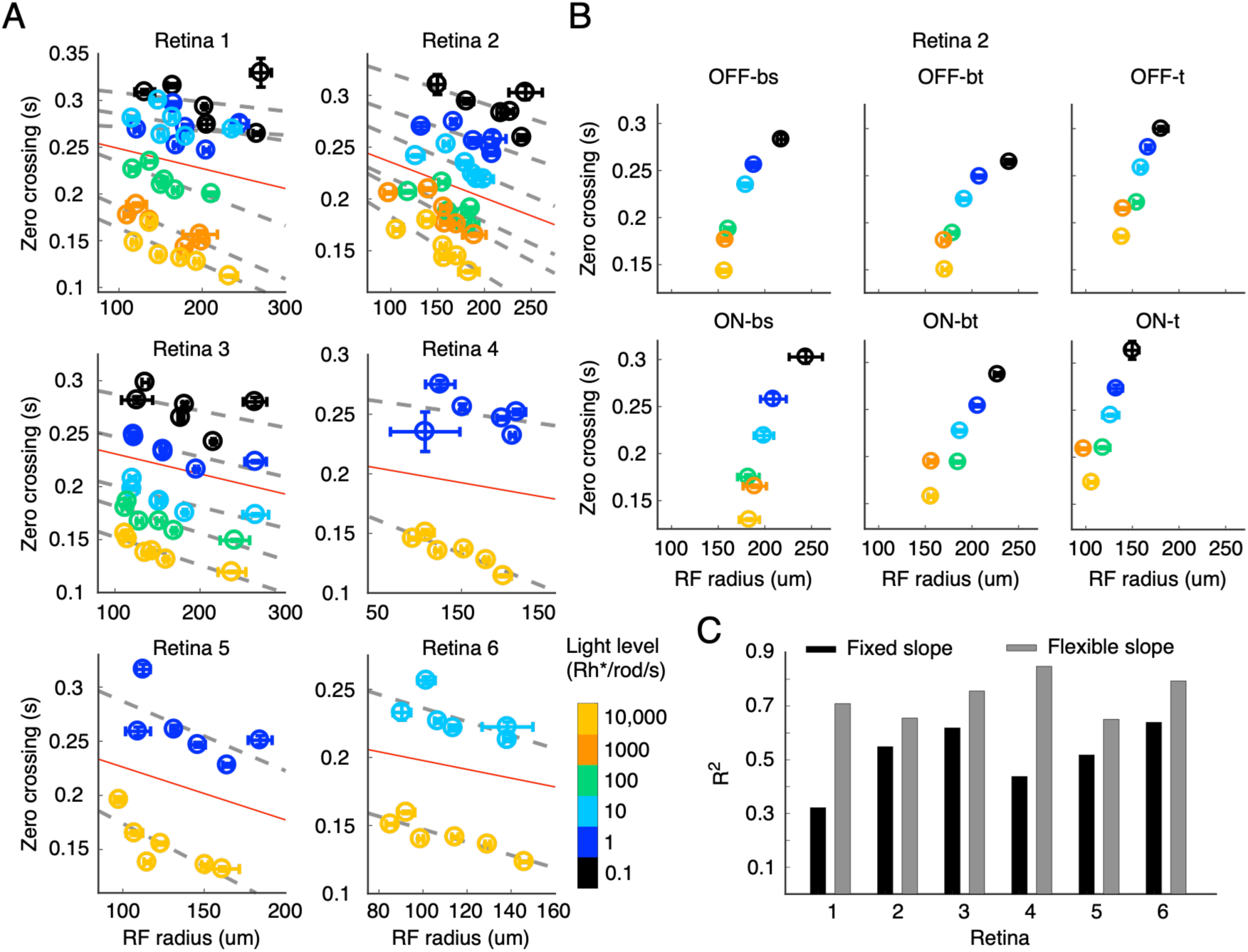
**Space-time tradeoff is present at all light conditions**. (**A**) The relationship between spatial and temporal RF resolution for each type and light level in six retinas. Increased spatial resolution, or smaller RF centers, correlates with decreased temporal resolution, or longer temporal integration. Data points are averages with SEM. (**B**) Plotting space-time tradeoff for each cell type, using data from the 2nd retina of **A**. (**C**) Performance of a fixed-slope best fit line (red lines in **A**) compared to a best fit line with flexible slopes over light levels (dashed lines in **A**. The total number of RGCs across six retinas depended on light level. For OFF-bs, the range of recorded RGCs over all light levels is 34-159; OFF-bt: 48-237; OFF-t: 10-82; ON-bs: 20-110; ON-bt: 52-175; ON-t: 16-83.

### Relative contrast gain is stable with light adaptation

Spatial and temporal RFs identify the linear component of spatial and temporal integration in RGCs. However, they do not describe how stimuli of increasing spatiotemporal contrast change spiking responses of RGCs. Thus, we next analyzed the stability of the contrast gain function of these six RGC types across light levels. In the framework of a linear-nonlinear-Poisson model, each RF is the linear stage, and the contrast response function is the static nonlinear component of the model. The contrast gain function summarizes the relationship between the (RF filtered) stimulus and spike rate, which can be more or less rectifying (Chichilnisky 2001; Chichilnisky and Kalmar 2002). We found that the static nonlinearities across cell types are remarkably stable over light levels from scotopic to photopic conditions (**Figure 9**). To quantify the shape of these static nonlinearities, we used a nonlinearity index (NLI) in which more negative values signify a linear relationship between contrast and spike output and more positive values indicate strongly rectified, nonlinear gain (Chichilnisky and Kalmar 2002). NLIs were systematically related to the temporal filtering properties of the RGCs: OFF-bs, ON-bs, and OFF-bt were the least nonlinear, while OFF-t, ON-t, and ON-bt show ed the most rectification. These relationships were consistent across light levels (p<0.05). There was some significant reordering of these relationships at 1000 Rh*/rod/s (**Figure 9B**). This is very near the light level where cone signals begin to dominant rod signals (Naarendorp et al. 2010) and may therefore be produced by in an imperfect handoff from rod to cone signaling. Nevertheless, the relative organization of contrast response functions across RGC types is remarkably well conserved from scotopic to photopic conditions.

**Figure 9:**
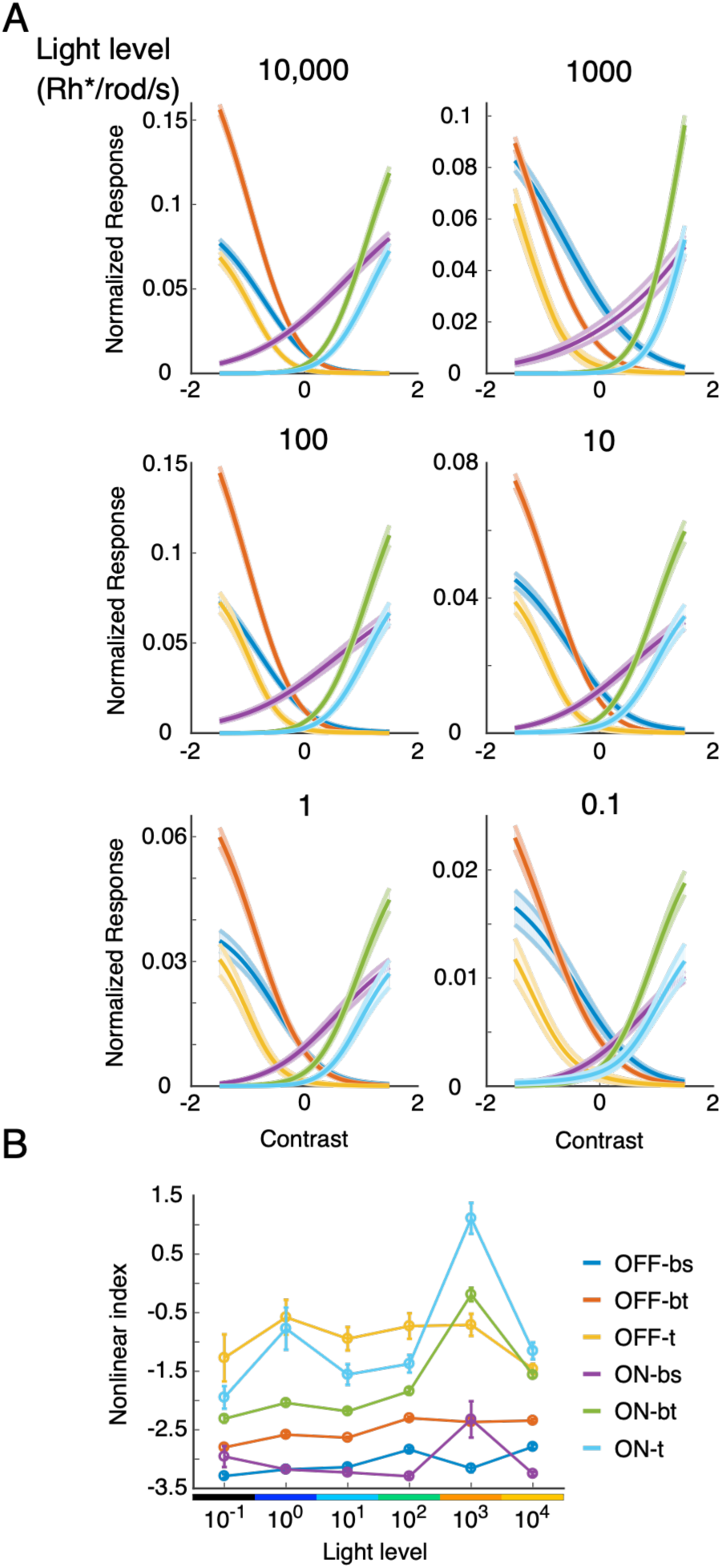
Contrast response functions across RGC types maintain stable organization from scotopic to photopic conditions. (**A**) Average SNLs over four retinas with SEM as shaded regions. (**B**) Nonlinearity index across light levels. The brisk sustained types are the most linear, the brisk transient are in the middle, and the transient types are the most nonlinear. This organization is generally consistent across light levels. The total number of RGCs depended on light level. For OFF-bs, the range of recorded RGCs over all light levels is 35-107; OFF-bt: 48- 160; OFF-t 16-63; ON-bs 24-51; ON-bt 52-130; ON-t 21-62.

### Persistence of band-pass tuning at low light levels

Initial investigations of efficient coding theory applied to natural scene statistics predicted that spatial and temporal RFs should transition from bandpass at high signal-to-noise ratios (SNRs) to low-pass at low SNR (Atick and Redlich 1990, 1992). However, these studies made several simplifying assumptions, such as assuming a linear input-output relationship for the encoding units (Atick and Redlich 1992; van Hateren 1992; Srinivasan, Laughlin, and Dubs 1982). More recently, an alternative linear-nonlinear model of retinal encoding has been developed that implements efficient coding by directly optimizing mutual information between system inputs and outputs (**Figure 10A-B**) (Karklin and Simoncelli 2011; S. Roy et al. 2021; Jun, Field, and Pearson 2021, 2022). This ‘Karklin and Simoncelli’ formulation predicts the presence of ON and OFF pathways with center-surround RFs, and multiple cell types. Therefore, we investigated what the Karklin and Simoncelli model predicts about center-surround RF structure across a range of low-to-high SNR conditions. Surprisingly, we found that this linear-nonlinear model did not predict a loss of RF surrounds with increases in input noise (analogous to decreasing the light level). Instead, the radii of both the center and the surround increased with increasing input noise (**Figure 10C**).

**Figure 10:**
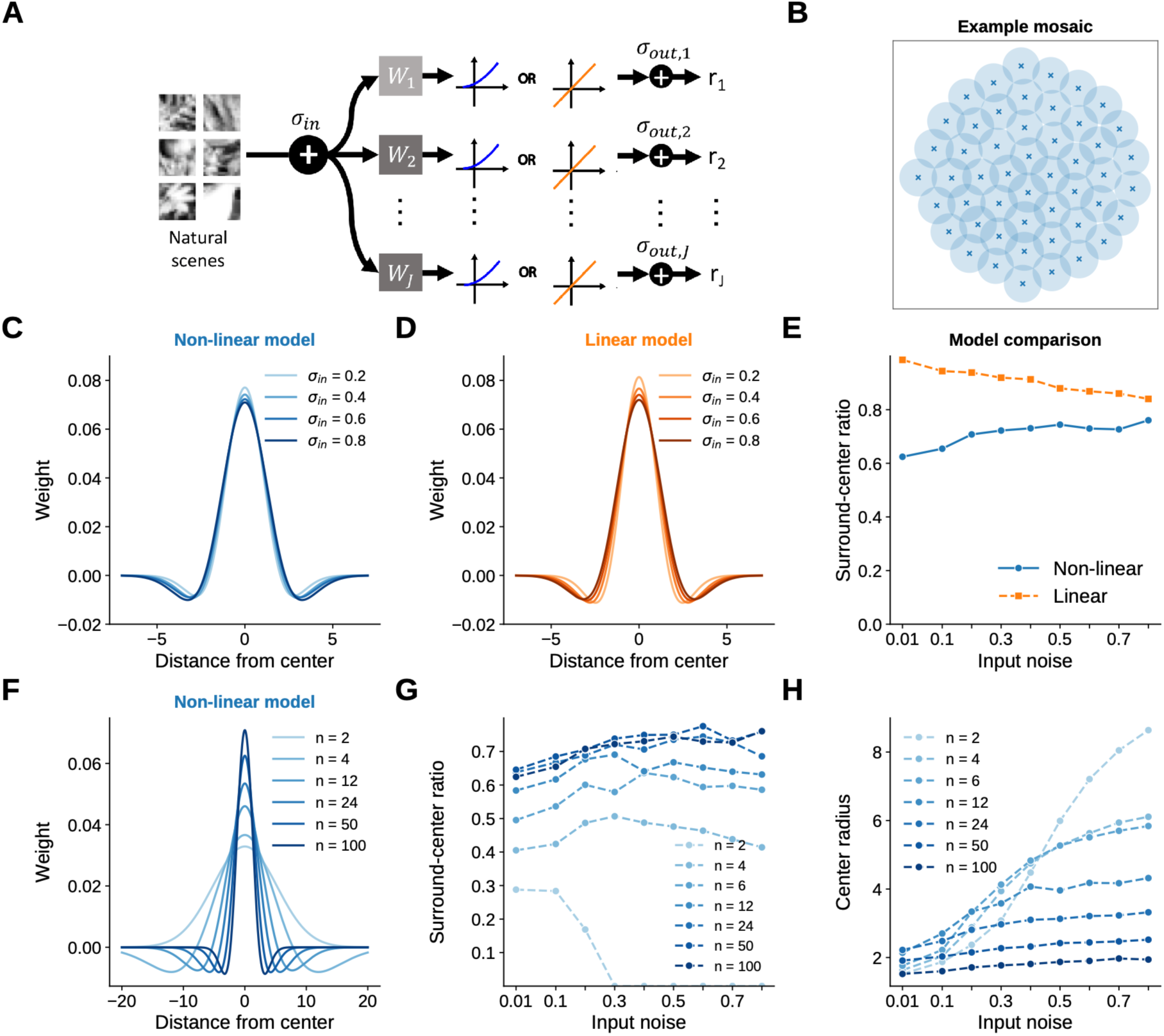
Efficient coding predicts center-surround (band-pass) tuning across a range of signal-to-noise values. (**A**) Model schematic: natural scenes are passed through a set of spatial kernels (*w_j_*) that mimic RFs; the outputs are then passed to a static nonlinearity (blue) or a static linear function (orange). These yield instantaneous firing rates (*r_j_*) for each model neuron (unit) and image. Noise is added at the input (*σ_in_*) and output (*σ_out_*) of the model. (**B**) RFs of 50 units resulting from optimizing the model in **A** to maximize the mutual information between a library of natural images and the output firing rates. Each circle summarizes the RF of one unit and is drawn at 0.35 of the maximum RF value. The RFs tile space. (**C**) The mean RF shape of 100 units in the nonlinear model for a range of input noise values: note the RFs retain a surround at high noise values. (**D**) Same as (**C**) but for the linear model with 50 units. Halving the unit number is justified because the nonlinear model yields ON and OFF units, while the linear model produces one mosaic that can signal both positive and negative firing rates. (**E**) Ratio of the surround to center RF areas plotted as a function for input noise for both models. (**F**) Mean RF shapes as the number of units in the nonlinear model is varied. At small unit numbers, the surround disappears as the RFs have more room to expand. (**G**) Ratio of center to surround strength plotted as a function of input noise for models with different numbers of units. Only at small unit numbers does increasing input noise lead to a weaker RF surround. (**H**) Center radius as a function of input noise for different numbers of units in the model. Increasing input noise increases the RF radius.

We wondered if this discrepancy with previous results (Srinivasan, Laughlin, and Dubs 1982; Atick and Redlich 1990, 1992; van Hateren 1992) was caused by the presence of the rectifying nonlinear output; prior work assumed linear units. To test this possibility, we imposed linear output functions and reoptimized the model across a range of input noise values (**Figure. 10A**, orange). Again, the resulting receptive fields retained a surround but increased the radius of both center and surround (**Figure 10D**). The surround persisted despite matching SNR values to those of previous studies (Atick and Redlich 1992).

In summary, both linear and nonlinear output functions redict a weak dependence of RF surrounds on input noise. The linear model did tend to yield somewhat stronger RF surrounds than the nonlinear model (**Figure 10E**). However, these results indicate that the difference between predictions previous formulations and the Karklin and Simoncelli formulation is not caused by the inclusion of nonlinear outputs.

Another difference between the two formulations is that previous studies assume translation invariance: every ganglion cell kernel is the same function translated across an infinite retina, so the population objective collapses to that of a single cell. This assumption leaves the density of the population unspecified. The Karklin and Simoncelli formulation instead places a specified number of units over a finite image, so the spacing between neighboring RFs is determined and the optimization must minimize any redundancy that spacing produces. To test if this difference is consequential, we reduced the number of units in the Karklin and Simoncelli model to mimic a reduction in neighbor-neighbor interactions. As unit number was reduced and noise increased, we found that RF surrounds disappeared at high noise values (**Figure 10F-G**). These results indicate that spatial interactions between well-packed RFs cause a persistence in RF surrounds even under high noise (low SNR) conditions. Increasing input noise did results in larger RF radii regardless of the number of units in the model (**Figure 10H**). In conclusion, the observation of persistent RF surrounds in rat RGCs at low light levels (**Figure 7**) is consistent with efficient coding theory when the analysis includes neighboring RFs spatially interacting over a finite viewing area.

## Discussion

How does light adaptation affect parallel processing in the retina? We have laid out several possibilities (**Figure 1**), all of which have support in the literature. To distinguish between these options for a well-defined population of six RGC types -- those with center-surround RFs that are approximately space-time separable -- we simultaneously measured their RF structure across a million-fold change in light level. We found that the impact of light adaptation on parallel processing was broadly consistent with proportional adaptation, where the relationships among different cell types are preserved between sunlight and starlight. Temporal integration in ON-bs RGCs and spatial integration in ON-bt RGCs were exceptions to this trend, showing unique RF changes that result in mixed adaptation. We did not find much evidence for less functional diversity across cell types (inhomogeneous adaptation) or a large-scale reorganization of visual signals, both suggested by some previous studies (Jakiela and Enroth-Cugell 1976; Tikidji-Hamburyan et al. 2015). The observation that proportional adaptation is predominant suggests that visual features encoded by the *relative* responses across different RGC types may provide invariant signals to the brain from starlight to daylight. Below we compare these results to previous experimental and theoretical studies, speculate on mechanisms that could contribute to ‘mixed’ adaptation, and discuss potential implications for downstream visual processing.

### Multiple possible forms of population adaptation

This study is distinct from previous work because it focuses on understanding how light adaptation alters the relative RF properties across many identified and irreducible RGC types. The types are ‘identified’ because a rigorous functional classification defined each type (Ravi et al. 2018). They are ‘irreducible’ because each type forms a mosaic and thus cannot be further subclassified (Wassle, Peichl, and Boycott 1981; Devries and Baylor 1997; Field et al. 2007). In general, light adaptation could impact parallel processing in several ways, including proportional, inhomogeneous, reorganizing, or mixed adaptation (**Figure 1**). Notice that conceptually, adaptation could have different population-level effects on spatial versus temporal RFs. For example, spatial RFs could adapt proportionally, while temporal RF adaptation could be inhomogeneous (Jakiela and Enroth-Cugell 1976), and the effect on contrast gain could be reorganizing, such as a switch from ON to OFF responses (Tikidji-Hamburyan et al. 2015). Thus, there is a rich diversity of ways adaptation could alter parallel processing. Our results are consistent with ‘mixed’ adaptation, but this mixed adaptation was primarily composed of ‘proportional’ adaptation. Except for the temporal and spatial RFs of ON-bs and ON-bt RGCs, respectively, the impact of light adaptation on visual processing largely maintained the organization of the relative relationships across cell types. This preservation of relative signaling across types was particularly striking for contrast gain (**Figure 9**), in which the relative signaling was remarkably stable across a million-fold light intensity range spanning rod to cone vision.

As noted above, ON-bs RFs represent one counterpoint to proportional adaptation. ON-bs temporal integration sped much more from scotopic to photopic conditions than any other type (**Figure 4B-C** and **5B**). This cell type also exhibited a profound change in temporal filtering, switching from low-pass under scotopic conditions to band-pass under photopic conditions (**Figure 4F** and **5A**). This observation suggests that not all the changes in temporal integration across RGC types and light levels could be accounted for by changes in photoreceptor kinetics. One possible and somewhat speculative explanation for the outstanding nature of ON-bs RF adaptation is the potential expression of melanopsin by these neurons. ON-bs RGCs likely correspond to the morphologically defined ON-sustained alpha cells (Krieger et al. 2017), which in the mouse retina express melanopsin (Estevez et al. 2012; Sexton et al. 2015). Melanopsin activation has been shown to impact contrast coding (Sonoda et al. 2018; Daly et al. 2026), and it is conceivable that changing the resting membrane potential of the ON-bs RGCs across light levels affects their temporal adaptation differently from RGCs that lack melanopsin. In addition, ON- bt RGCs exhibited a clear loss of the RF surround at the lowest light level, while the other RGC types retained a strong surround. We do not know why the ON-bt RGCs lost their RF surround, though it is suggestive that the surround is primarily generated post-synaptic to AII amacrine cells under these conditions (Bloomfield and Dacheux 2001).

This study also differs from previous work in analyzing the effects of adaptation on multiple RGC types that are irreducible (i.e., they cannot be further subclassified). Each of the six RGC types we examined exhibited strong evidence for a mosaic-like organization (**Figure 1**) (Ravi et al. 2018). Previous studies of adaptation that include multiple cell types either rely on cell type classification with histological markers that do not uniquely identify irreducible cell types (Farrow et al. 2013; Tikidji-Hamburyan et al. 2015), or simply separate RGCs by their ON and OFF polarity (Cowan et al. 2017; Borghuis, Ratliff, and Smith 2018). Observing mosaics was made possible by using large-scale multi-electrode arrays, which also allowed tracking individual cells across a wide range of light levels using electrical images (Field et al. 2009; Yao et al. 2018). These technical differences, and a focus on six RGC types that exhibit relatively simple RF structures, may explain why we did not observe a large-scale reorganization of retinal signaling suggested by some previous work (Tikidji-Hamburyan et al. 2015). Indeed, at least some of the preferred polarity changes across light levels are due to direction-selective RGCs (Pearson and Kerschensteiner 2015) that we did not examine in this study.

### Comparison to previous literature on low-pass filtering at low light levels

The canonical view of light adaptation is that spatial RFs expand and temporal RFs are prolonged at low light levels. Our measurements are consistent with this view. However, the canonical view also specifies that at high light levels, RGCs differentiate visual input to remove spatial and temporal correlations present in natural scenes. However, at lower light levels, they switch to purely integrating visual input to mitigate the consequence of noise (Atick and Redlich 1992; van Hateren 1992). In space, differentiation takes the form of an antagonistic relationship between the center and surround. In time, differentiation takes the form of a biphasic temporal RF. In the frequency domain, this switch from differentiation to integration corresponds to band-pass and low-pass filtering, respectively. Some studies agree with these predictions, finding that RFs shift to low-pass filtering in space and time at low light levels (Barlow, Fitzhugh, and Kuffler 1957; Muller and Dacheux 1997; Enroth-Cugell and Shapley 1973; Jakiela and Enroth-Cugell 1976; Purpura et al. 1990; Farrow et al. 2013; Borghuis, Ratliff, and Smith 2018). Importantly, several other studies have produced an alternative view that at least spatial frequency tuning remains band-pass under scotopic conditions (Troy, Oh, and Enroth-Cugell 1993; Troy, Bohnsack, and Diller 1999; Tikidji-Hamburyan et al. 2015; Cowan et al. 2017). Complicating the interpretation of these findings is the fact that many of these studies focused on different RGC types in various species, including mice, rabbits, cats, and primates.

Based on our measurements of hundreds of RGCs recorded simultaneously from six identified and irreducible types, we observe that most RGC types in rat maintain robust band-pass spatial and temporal filtering under scotopic conditions as low as 0.1 Rh*/rod/s. Only two RGC types -- ON-bs and ON-bt -- transitioned to more low-pass filtering in the temporal and spatial domains, respectively. Evidence for a transition from band-pass to low-pass spatial tuning in at least some cat and primate RGC types is compelling (but see ((Troy, Oh, and Enroth-Cugell 1993; Troy, Bohnsack, and Diller 1999)), suggesting that the rat, and possibly the mouse, retina operates under somewhat different signal and noise constraints than cat and primate retinae. Still, some previous studies have suggested a switch to low-pass filtering at low light levels in the rodent retina (Farrow et al. 2013; Borghuis, Ratliff, and Smith 2018). A possible explanation is that the stimulus intensities we used were not low enough to elicit spatial or temporal filtering changes. However, this seems unlikely as we utilized light levels 100-fold dimmer than those studies. Furthermore, at light levels lower than 0.1 Rh*/rod/s, Poisson variability in photon absorptions becomes large relative to the contrast of a drifting grating (or checkerboard stimulus), making it challenging to measure RF properties at these ultra-low scotopic conditions. Overall, our findings show that even in starlight conditions, many rat RGCs perform band-pass filtering in space and time, necessitating a more nuanced view of the effects of light adaptation on RF structure, at least in the rodent retina.

Some prior work has indicated an abrupt switch in RF structure between rod and cone mediated light levels (Farrow et al. 2013), while other studies have indicated a smoother transition (Y. V. Wang, Weick, and Demb 2011). We observe some instability in the contrast response functions in the mesopic range (**Figure 9B**, ∼1000 Rh*/rod/s), but generally the transition in RF structure from rod to cone light levels is relatively smooth. However, it is possible that with a finer sampling of light levels, we would observe a discontinuity in the transition between the rod and cone dominated signaling.

### Implications for efficient coding theory

The persistence of a RF surround at low light levels appears to be at odds with previous predictions made by efficient coding theory, which is thought to prescribe a center-surround RF at high light levels (a high signal-to-noise regime) and a center-only RF at low light levels (a low signal-to-noise regime) (Atick and Redlich 1992; van Hateren 1992; Srinivasan, Laughlin, and Dubs 1982). However, previous these predictions assume a linear system and RGCs exhibit a range of nonlinearities, including rectification in their spike output (**Figure 9**). More consequentially, previous work effectively optimizes a single output unit and does not specify an RGC density over space. Atick and Redlich (1990, 1992) assume translation invariance, so every ganglion cell kernel is a translated copy of every other across an effectively infinite retina, and the objective reduces to that of one cell. They do penalize redundancy between neighboring cells, since decorrelating the outputs of the ganglion cell array is their objective. However, what their solution specifies is a correlation length between those outputs rather than the redundancy of any particular array of RFs. Whether that correlation length implies redundancy among real neighbors depends on how many cells cover a given area, which their formulation never specifies. Srinivasan and colleagues (1982) similarly derive the surround of a single neuron given a grid of input units, rather than a grid of output neurons that jointly sample space. Finally, the van Hateren (1992) study is exclusively in the frequency domain. Therefore, none of the previous work on RF structure and its dependence on the SNR of sensory inputs fixes the number of output units over a given area. Thus, the redundancy present among neighboring RFs is never a determined quantity in these formulations.

We investigated the predictions of efficient coding theory when applied to a more accurate (though still incomplete) model of retinal output: a fixed set of linear-nonlinear units (**Figure 10**) optimized on real images, not just their second-order correlation structure (Karklin and Simoncelli 2011; Jun, Field, and Pearson 2021). We found that under this Karklin and Simoncelli formulation, center-surround structure persisted over a large (100-fold) range of input noise values provided there were more than a few RFs spanning the images (**Figure 10**). Qualitatively, this result did not depend on whether the output was linear or nonlinear. However, when only a couple of RFs were optimized in the nonlinear model, the surround strength did depend strongly on the amount of input noise, with higher levels of input noise resulting in a diminished surround (**Figure 10G**).

We interpret this result as indicating that under efficient coding theory, when RFs are not forced to interact with their neighbors, coding efficiency pushes them to increase the extent of their spatial integration (center size) and lose their spatial antagonism. However, when the neighboring RFs are tightly packed, losing spatial antagonism results in unacceptably high levels of redundancy even under low SNR conditions: the “pressure” to expand is counteracted by a reciprocal “pressure” from nearby RFs. To combat the effects of increased input noise, the RFs instead increase the spatial integration of both the center and surround, while keeping the ratio of their integrated volumes approximately constant (**Figure 10**). This prediction is qualitatively consistent with how the majority of the RGC types adapted to decreasing light levels in this study (**Figure 7**).

We also note that not all the RGC types strictly obeyed this prediction. In particular, the ON-bt RGCs displayed the classical behavior of becoming center-only at the lowest light levels tested. Importantly, a full exploration of the predictions of efficient coding theory, including a comparison to our observation that temporal RFs remain mostly bandpass across light levels, will require optimizing more sophisticated models of the retina that include space, time, chromatic channels and multiple nonlinearities (Jun, Field, and Pearson 2022; Brinkman et al. 2016). It is possible that once models are sufficiently complex to predict multiple cell types (e.g., (Jun, Field, and Pearson 2022)), they may also predict some heterogeneity in how those cell types adapt to changes in stimulus statistics.

### Caveats and limitations

One potential limitation of our results is relating in vivo dynamics to our ex vivo measurements. The isolated retina preparation used here removes the recycling of chromophore by the pigment epithelium. At 100 and 1000 Rh*/rod/s, the amount of bleached chromophore that builds up in rod photoreceptors may depart from equilibrium chromophore recycling when the epithelium is present (Lamb and Pugh 2004; Borghuis, Ratliff, and Smith 2018). These bleaching-induced changes in rod sensitivity are likely to impact RGC spiking, though it is unclear the extent to which they might influence spatial or temporal RF structure. However, for dimmer stimulus intensities (<100 Rh*/rod/s), rod bleaching is not a significant factor because <1% of the pigment is bleached over the duration of these measurements (Leibrock, Reuter, and Lamb 1998). Similarly, at 10,000 Rh*/rod/s, cones are bleached at a relatively slow rate: ∼10% of photopigment per hour, assuming no pigment regeneration from residual pigment granules on the retina or Muller glia (J. S. Wang et al. 2009). In addition, cones frequently operate under conditions with substantial amounts of chromophore bleaching in vivo (Burkhardt 1994; Valeton and van Norren 1983). A conservative interpretation of our data would be only to consider results at 10,000 Rh*/rod/s and <100 Rh*/rod/s as matching in vivo conditions. Even from this perspective, we can conclude that RF organization across different RGC types is dominated by proportional adaptation at low scotopic to low photopic conditions, with potential deviations in the mesopic range.

The *ex vivo* conditions of these experiments necessitated examining adaptation from low light levels to high light levels. Without the pigment epithelium attached to the retina, the rods cannot dark adapt. Thus, animals were dark-adapted overnight prior to initiating the experiments, then euthanasia and dissections were performed in complete darkness with the assistance of infrared converters. This caused the retina to be in a high-sensitivity state at the beginning of the experiment. It is possible that this experimental design impacted the results. For example, the presence of a strong surround in many RGC types at the lowest light level could have resulted from allowing a long time for dark adaptation to reach steady state.

Additionally, isolating the retina likely perturbs the *in vivo* concentrations of neuromodulators that are related to circadian rhythms and light adaptation, particularly dopamine and melatonin. These neuromodulators play important roles in light adaptation by, for example, alternating the extent of gap junction coupling at numerous cites across the retina (Suva Roy and Field 2019; Bhoi et al. 2023). The extent to which these processes are impaired in our ex vivo experiments is not clear. Thus, an important direction for future work is to understand how these neuromodulators impact receptive field structure and contrast response functions.

Another caveat is that we examined the effects of light adaptation by assaying changes to the linear component of the RF, which misses some aspects of RF structure. For example, a linear-nonlinear-Poisson (LNP) model describes 30-70% of the response variance in RGC responses in our experiments (**Figure 2 -supplemental figure 1**). RF mechanisms that account for the rest of this variance may also change across light levels. For instance, rectified spatial integration among subunits (Hochstein and Shapley 1976), which the LNP model cannot capture, has been shown to decrease with light level in ON alpha RGCs (Grimes, Schwartz, and Rieke 2014). We emphasize, however, that the RFs we analyzed capture first-order response changes in RGCs and thus are a first step in understanding how parallel processing of diverse cell types is altered by light adaptation.

Finally, our observations were made using artificial visual stimuli. Previous work has shown that some RF properties are specific to naturalistic stimulation. For example, primate OFF parasol RGCs linearly integrate spatial patterns of artificial stimuli but exhibit nonlinear responses when stimulated with patches of natural scenes (Turner and Rieke 2016; Turner, Schwartz, and Rieke 2018). Other work has similarly shown that center-surround interactions can exhibit nonlinearities under natural scenes (Takeshita and Gollisch 2014). Thus, it will be important for future work to determine how the encoding of naturalistic stimuli interacts with light adaptation and parallel processing across many RGC types.

### Implications for downstream processing

The same visual stimulus presented at different light levels produces remarkably different spiking patterns in individual neurons (Tikidji-Hamburyan et al. 2015). This is a direct result of RF structure changing across light levels. However, these response changes challenge downstream circuits to achieve stable processing of visual input across light levels. We suggest that the proportional adaptation across many RGC types shown here may provide a potential solution, where an invariant signal results from comparing the relative responses between different RGC types. This coding strategy would only apply to specific combinations of cell types and could constrain the downstream convergence of signals to only those cell types exhibiting proportional adaptation. This possibility is consistent with previous findings about cell type-specific projections from the retina to the LGN. Some neurons receive input from several cell types, so extracting invariant features may be a useful motif for these cells (Rompani et al. 2017; Liang et al. 2018; Roman Roson et al. 2019). Other LGN neurons receive input from mainly one RGC type; these would not be able to process relative responses of different types. Instead, those LGN neurons may either pass along light adaptation changes to other regions or shift their processing of retinal activity with light level.

Few studies have examined the impact of mean light intensity in post-retinal brain areas. One study measured responses in mouse LGN across light levels, finding that response polarity changes are inherited from the retina (Tikidji-Hamburyan et al. 2015). Other studies in cat and primate LGN and V1 indicate small changes in RFs across light levels (Virsu, Lee, and Creutzfeldt 1977; Bisti et al. 1977; Wiesel and Hubel 1966; Ramoa, Freeman, and Macy 1985; Duffy and Hubel 2007; Rhim, Coello-Reyes, and Nauhaus 2021). These studies largely found stability in response properties such as band-pass filtering, orientation selectivity, and direction selectivity over light levels. Understanding the extent to which V1 and other visual areas produce invariant responses across light levels and how this is achieved given the changes in feature selectivity in the retina remains an area open to future investigation.

## Materials and Methods

### Multi-electrode array recordings

All experiments were performed in accordance with the guidelines of the Institutional Animal Care and Use Committee (protocol #A200-19-00). Long-Evans rats (9 females, 3 males; ages 60-250 days; Charles River Laboratories) were dark-adapted overnight and euthanized with an intraperitoneal injection of ketamine/xylazine followed by decapitation. Euthanasia and retinal dissections were performed in darkness with the assistance of infrared converters. Approximately 3 mm x 2 mm dorsal pieces of retina were dissected and placed RGC-side down on an electrode array. Two MEAs were used: one had 512 electrodes with 60 µm spacing covering an area of 0.9 mm x 1.8 mm, while the other consisted of 519 electrodes with 30 µm spacing covering a hexagonal area that was 0.48 µm across (Litke et al. 2004; Frechette et al. 2005; Field et al. 2010). The tissue was perfused with oxygenated Ames solution at a rate of 6-8 mL/min and passed through an inline solution heater set to 34°C. The voltage on each electrode was sampled at 20 kHz and filtered to preserve frequencies between 80 and 2000 Hz.

### Spike sorting and neuron identification

Spikes on each electrode were identified by thresholding and selecting voltage deflections that exceeded four times a robust estimate of the standard deviation of voltages across time. Spikes were sorted and clustered to identify putative neurons (RGCs and spiking amacrine cells) in three steps: 1) dimension reduction by principal components analysis; 2) clustering initialization by a water-filling algorithm; 3) fitting a mixture of Gaussians model to the data by expectation maximization (Shlens et al. 2006; Field et al. 2007). Clustered spikes were identified as neurons only if they exhibited a refractory period (1.5ms) with <10% estimated contamination and were composed of at least 100 spikes. To track identified RGCs across light conditions, cell clusters were sorted in the same PCA subspace at each light level. Neuron identity was verified by checking that electrical images (EIs) and RF locations were stable across conditions (Field et al. 2009; Yao et al. 2018).

### Visual stimuli, receptive field measurements and RGC classification

Visual stimuli were created with custom Matlab code. Stimuli were presented with a gamma-corrected OLED display (SVGA+XL Rev3, Emagin, Santa Clara, CA). The image from the display was focused onto the photoreceptors using an inverted microscope (Ti-E, Nikon Instruments) with a 4x objective (CFI Super Fluor 4x, Nikon Instruments). The optimal focus was confirmed by presenting a high spatial resolution checkerboard stimulus (20 µm x 20 µm checkers, refreshing at 15 Hz) and adjusting the focal plane of the stimulus to maximize the average spike rate over all electrodes. The intensity of the stimulus was set using neutral density filters in the light path. In each experiment, stimuli were presented in increasing order of light intensity. The tissue was adapted at the next higher light level for at least ten minutes before measuring responses.

Checkerboard noise was used to measure spatial RFs, temporal RFs, and contrast response functions by computing the spike triggered average and an associated static nonlinearity that describes the relationship between the filtered stimulus and the instantaneous spike rate (Chichilnisky 2001; Chichilnisky and Kalmar 2002; Ravi et al. 2018). All checkerboard noise stimuli were black-white, meaning the red, green and blue primaries of the video display took the same value (0.1 or 0.99) within a checker. The checkerboard contrast was always 98% Michelson contrast. In most experiments, the size and frame rate of the checkerboard noise were adjusted at each light level to account for the changes in the spatial and temporal integration performed by the RGCs. At the lowest light level, 0.1 Rh*/rod/s, the largest checker size used was 252 µm x 252 µm and the slowest refresh was 15 Hz. At the highest light level, 10,000 Rh*/rod/s, the smallest checker size used was 63 µm x 63 µm and the fastest refresh was 60 Hz. In control experiments, these parameters were varied at a single light level and the estimated RF properties were compared for different checkerboard stimuli. Within the set of parameters we used, RF estimates were indistinguishable up to a constant scaling factor for different spatial and temporal parameters (**Figure 4 -supplemental figure 1**). Stimuli consisted of non-repeated, binary noise patterns interleaved with repeated, binary noise sequences (5 or 10s in duration) to control for non-stationarities in recordings and validate model fitting.

Spatial RFs were defined by the stimulus pixels in the STA with intensity values that exceeded four standard deviations from the mean. Temporal RFs were computed by averaging the intensity profile of the pixels composing the spatial RFs across time in the STA. The transience index was computed by taking the difference between the areas under the first and second lobes on the temporal RF and dividing by their sum (**Figure 3F**). RFs with low signal-to-noise ratios were excluded from analysis.

Drifting sinusoidal gratings were used to measure the spatial frequency tuning functions of RGCs across light levels (Enroth-Cugell and Robson 1966). The grating comprised 11 randomly interleaved spatial frequencies with periods spanning 210 to 6720 µm (or ∼7 to 220 degrees of visual angle). Their temporal frequency was 2Hz and the contrast was set to 50% except at 0.1 Rh*/rod/s where the contrast was 100%. In each experiment, results were replicated for gratings with contrasts set to 25% (50% for 0.1 Rh*/rod/s). The higher contrasts were used at 0.1 Rh*/rod/s to reduce the impact of Poisson variability in photon absorptions in estimating the responses at the lowest light levels. Some RGCs exhibited particularly noisy spatial frequency tuning curves that were not well fit by a difference-of-Gaussians. These cells were excluded from the analysis by the following criterion: we computed the root-mean-square deviation (RMSD) between the difference-of-Gaussians fit in the frequency domain and the normalized spatial frequency tuning curve for each cell. At each light level we calculated the median RMSD across cells and excluded cells with RMSDs larger than four median absolute deviations above the median. Depending on cell type and light level, between 8-14% of RGCs were excluded based on this criterion: the mean was 11.5% averaged across cell types and light levels.

To measure spatial RF sizes and the relative weighting between the center and surround from drifting grating stimuli, spatial tuning curves were generated by computing the power at the fundamental modulation frequency (2 Hz) for every spatial frequency. The tuning curves were then normalized to have a peak amplitude of one and fit with a difference of Gaussians function. This function consisted of four parameters representing the radius of the center, the radius of the surround and amplitudes for the center and surround. Parameters of the fit were estimated using fmincon in MATLAB with a global search and three constraints: the surround radius was restricted to be larger than the center radius, the surround radius was prevented from growing to sizes larger than ten-fold larger than the center radius, and the ratio between the area under the surround and that under the center was forced to be less than two. These constraints prevented the fitting function from using the surround function to capture the center and/or flipping the polarity of the cell’s predicted responses. The radii of the center and surround equaled the standard deviation of the fit Gaussians and the ratio between center and surround was calculated using the volumes of the two Gaussians. Note that some tuning curves did not level off with decreasing spatial frequency, so our estimates of surrounds are lower bounds.

Drifting square-wave grating were presented at the highest light level to identify direction-selective RGCs. These gratings were presented with spatial periods 126, 252, and 504 µm (or 4.2, 8.4, and 16.8 degrees of visual angle) at 1Hz temporal frequency and drifted in either 8 or 12 directions.

As described previously (Ravi et al. 2018), RGC types were classified at the photopic light level by first removing direction-selective RGCs and then clustering using the grating responses, RF properties, and autocorrelation functions computed from the spike times of each RGC. Classification was validated by observing a mosaic-like arrangement of spatial RFs for each identified type. Note that RGC spatial location was not included in the classification, and thus the mosaic arrangement was not imposed by the classification but instead reflected the underlying biological organization of irreducible RGC types in the mammalian retina.

### Space-time separability and tradeoffs

Space-time separability in the RFs was verified at different mean light levels by singular value decomposition (SVD). SVD factorizes the STA into pairs of spatial and temporal filters (**Figure 3 - supplemental figure 1**). Prior to performing SVD, the STA was spatially filtered by a Gaussian with a standard deviation of 0.75 times the checker size in the checkerboard stimulus. This substantially reduced noise in the STA while minimally perturbing the spatial structure. Consistent with previous results from these RGC types (Ravi et al. 2018), the space-time filter pair associated with the first singular value explained between 85% to 97% of the variance in the STA (**Figure 3B**). When the space-time filters associated with the second singular value exhibited some RF structure, this structure captured the temporal lag of the RF surround (**Figure 3 -figure supplement 1A**). Filter pairs associated with the third and higher singular values exhibited no clear structure related to the RF and appeared to only describe noise in the STA (**Figure 3 -figure supplement 1B)**.

The relationship between spatial and temporal resolution in RFs was quantified with two different linear fits of the data in **Figure 8**. First, we found a line that best fit all data points at a given light level, the so-called flexible fits. Next, to fit the data across all light levels, the spatial and temporal resolution were normalized by the average spatial and temporal resolution of all types at 10,000 Rh*/rod/s. We fit one line to these normalized points for a fixed fit and compared it to fitting lines separately at each light level (flexible fit). The performance of these fixed vs. flexible linear fits was quantified with R^2^.

### Contrast gain and linear-nonlinear analysis

Contrast gain functions were derived from the STAs and responses to the checkerboard noise as described previously (Chichilnisky 2001; Ravi et al. 2018). The checkerboard stimulus was filtered by the STA in space and time to produce a scalar value associated with each frame of the stimulus (the so-called generator signal). A histogram of spike counts for each generator signal point was produced and fit with a cumulative Gaussian function. The nonlinearity index (**Figure 9B**) was quantified as the log of the ratio of SNL slope at zero contrast to the slope at maximum contrast (Chichilnisky and Kalmar 2002). Outliers in nonlinearity index that exceeded 7 standard deviations beyond the mean were excluded.

### Experimental design and statistical analyses

The sample size was not predetermined by a statistical method, but sample sizes (number of RGCs cells and number of retinas) are similar to those generally used in studies of this kind. Experiments were replicated using at least three retinas and indicated in figure legends. Each retina was from a different animal. Statistical results are found in the text. Significance tests comparing different cell types were performed with the Kruskal-Wallis test (because data was generally not normal) using a Bonferroni correction for multiple comparisons. The Wilcoxon rank sum test was used for comparisons of a given cell type across light levels.

### Efficient coding models

A previously developed efficient coding framework for modeling retinal output was used to examine efficient coding predictions regarding the impact of input noise on RF structure **(Figure 10)** (Karklin and Simoncelli 2011; Jun, Field, and Pearson 2021). The model consists of an ensemble of simulated RGCs composed of a pixel weighting vector for each unit, *wj*, (modeling a receptive field) and a static nonlinear output function *η*(*y*_j_), (modeling contrast response functions): *j* denotes different RGCs, or units. Weight functions and output nonlinearities were optimized such that the mutual information between a library of natural images (Doi et al. 2003) and an ensemble of output spike rates was maximized. Independent and identically distributed (white) Gaussian input noise, *n_x_*, was added pixelwise to each input image (x). Gaussian output noise, *nr*, was added to the output firing rates. To compute the output firing rates, the inner product between the spatial filters *w*_j_ and noisy inputs, x + *n_x_*, was computed to get *y*_j_,

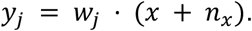

Where 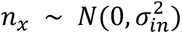 per pixel, and *N* is a normal distribution. The output of this dot product is then passed to the output nonlinearity to generate an instantaneous firing rate. The output nonlinearity was set as a softplus function,

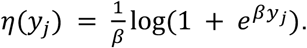

The β parameter was fixed to 2.5 to stabilize the optimization. The output firing rates, *r*_j_, were computed from η(*y*_j_) by using neuron-specific gain and threshold parameters *γ*_j_ and *θ*_j_, followed with independent additive noise 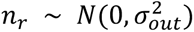,

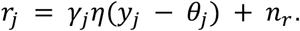

Note, in the linear version of this model (**Figure 10**) the output function was simply the identity,

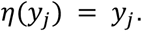

The optimized parameters of the model are the receptive field filter weights *w*_j_, the gain of the nonlinearity *γ*_j_, and the nonlinearity threshold *θ*_j_ of each output RGC. By assuming normally distributed inputs and locally linear responses, a closed-form approximation for mutual information can be derived, which leads to the following optimization objective (Karklin and Simoncelli 2011; Jun, Field, and Pearson 2021):

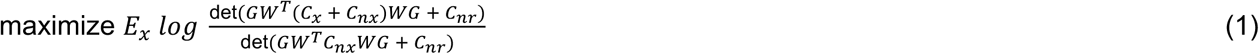

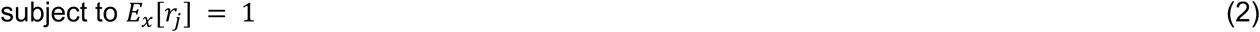

Here *C_x_* is the covariance matrix of *x*. *C_nx_* and *C_nr_* are the covariance matrices of input and output noises, respectively. The gain matrix *G* is diagonal, where its diagonal elements are the local derivatives of the output responses *r*_j_ with respect to *y*_j_ for each unit. In the nonlinear case, *G* increases as the firing rate increases, whereas it is independent of *x* in the linear case. In both cases, the training procedure involves estimating *G* across multiple samples and using this estimate to evaluate mutual information. *E_x_*[*r*_j_] is the average firing rate of each unit over images; this value is constrained to be one to impose a metabolic cost to spiking and was implemented via an augmented Lagrangian with a quadratic penalty (Nocedal and Wright 2006).

During training, *G* was estimated using a mini-batch of 128 image patches, which allowed calculating the gradients for *w*_j_, *γ*_j_, *θ*_j_. After taking a gradient step, the filters were renormalized to have unit norm. Models were trained for 1 million iterations, which was sufficient to ensure convergence of the mutual information. Unconstrained spatial kernels *w*_j_ have been shown to converge to characteristic center-surround shapes under optimization of mutual information (Karklin and Simoncelli 2011; Jun, Field, and Pearson 2021). Therefore, for increased speed and stability, these filters (RFs) were parameterized using a radially-symmetric difference of Gaussians:

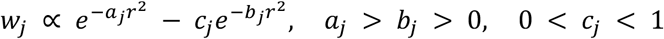

where *r* measures the spatial distance between a pixel and the RF center. The parameters *a*_j_, *b*_j_ and *c*_j_ determine the shape of each RF.

Parameters were optimized using adaptive moment estimation (Adam) (Kingma and Ba 2014) with a learning rate of 0.0001. Because *a*_j_, *b*_j_, *c*_j_, *γ*_j_ and *θ*_j_ need to be positive, the logarithms of the corresponding parameters were optimized. Also to speed the optimization and avoid local minima, jittering and centering were applied as described previously (Jun, Field, and Pearson 2021). Jittering was applied every 5,000 iterations between 20,000 and 100,000 iterations and consisted of moving the receptive field center by a random vector, with the x and y coordinates of this vector being drawn from a uniform distribution. Centering was applied between 100,000 and 400,000 iterations and consisted of adding an additional loss term that penalized large RFs and encouraged them to be localized around their centers. After these operations, the model was further optimized without jittering and centering for another 600,000 iterations. Jittering and centering were confirmed to not alter the final converged shape of the kernels nor any of the results in Figure 10. Also as described previously (Jun, Field, and Pearson 2021), a circular mask was applied to each input image (18x18 pixels) to minimize edge artifacts. The circular masking effectively leaves ∼255 image pixels in each input image patch x, and pixels outside the mask have input values near zero.

## Data and Code availability

Data and code to generate plots for figures 1-9 are included at the following GitHub repository: https://github.com/kmruda/RGC-functional-org. Data are also available in the Source Data files associated with each figure. Code for analyses associated with Figure 10 is available at https://github.com/pearsonlab/efficientcoding_adaptation.

## Acknowledgments

We thank Drs. Lindsey Glickfeld, Steven Lisberger and Jeffrey Beck for helpful discussion, and Ashley Zhou for assistance in spike sorting and pre-processing some data. This work was supported by R01 EY024567 (G.D.F.), R01 EY031396 (G.D.F.), and F31 EY028833 (K.R.).

## Author Contributions

K.R. and G.D.F. designed the study. K.R. and A.M.R perform electrophysiology experiments. K.R. and A.M.R analyzed the resulting data. N-Y.J., D.S-A. and J.P performed the theoretical predictions of efficient coding theory. K.R. and G.D.F wrote the manuscript. K.R., A.M.R, D.S-A. J.P., and G.D.F. edited the manuscript.

## Competing Interests

The authors have no competing interests.

---

**Figure 2-supplemental figure 1:**
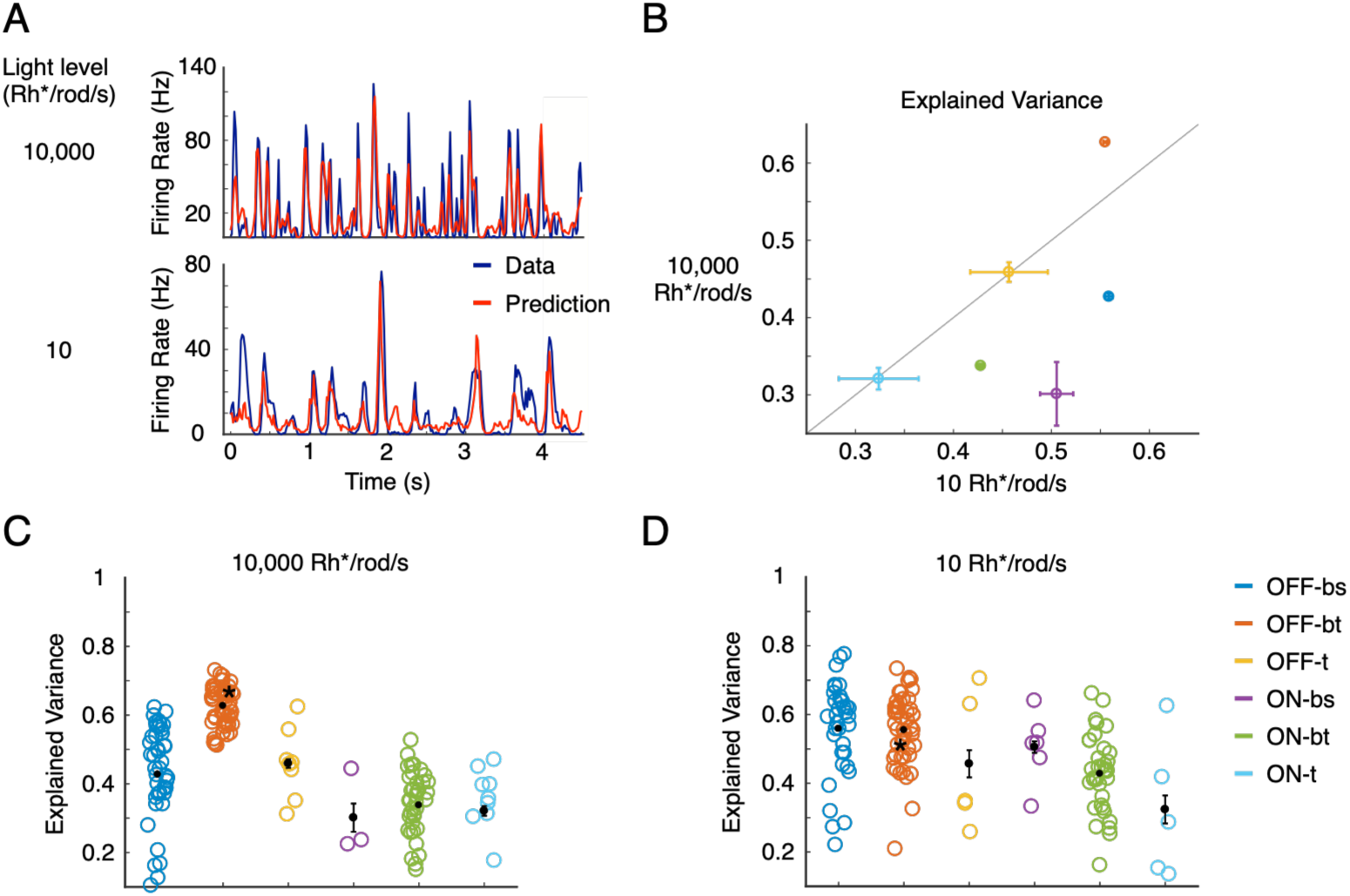
Linear-Nonlinear-Poisson (LNP) model predictions of RGC activity across light conditions. (A) Example RGC response to a white noise segment and the corresponding LNP prediction at a photopic (top) and scotopic (bottom) light level. (B) Comparison of LNP performance for the two light levels. Datapoints show mean with SEM bars and are from one retina. (C) Performance of the LNP prediction for all RGCs, broken down by cell type, at 10,000 Rh*/rod/s. The black asterisk marks the example RGC shown in A. (D) Same as C for responses at 10 Rh*/rod/s.

**Figure 3-supplemental figure 1:**
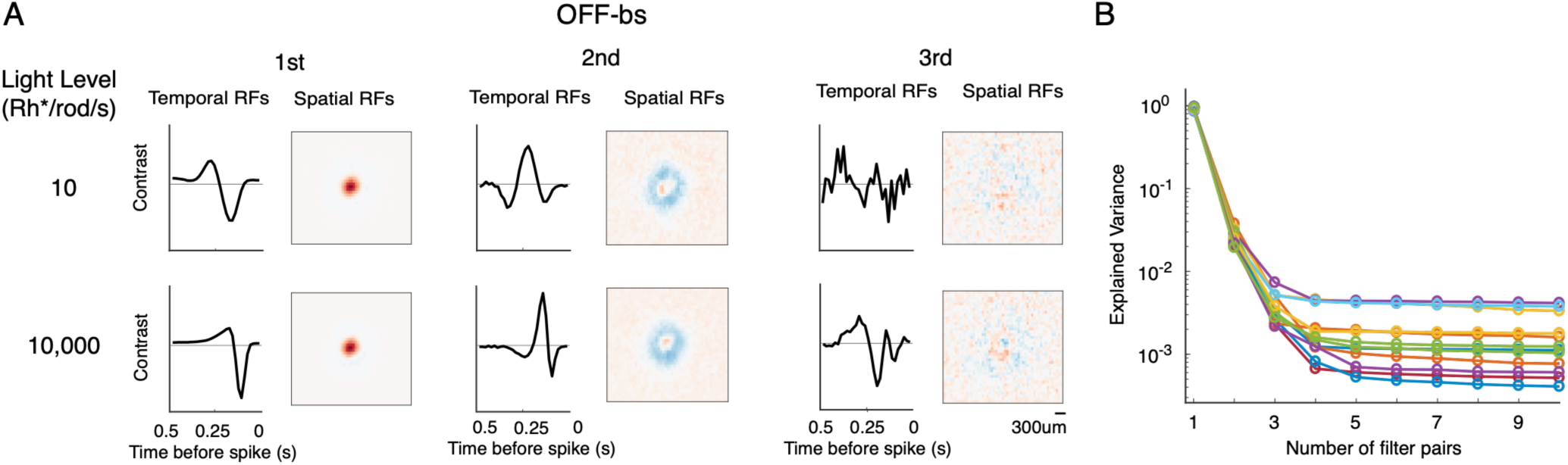
RF decomposition of RGC types. (A) Top 3 components of spatial and temporal filters for the average OFF-bs RF at a scotopic (top row) and photopic (bottom row) light level (data from one retina). If a RF is space-time separable, the first pair of spatial and temporal filters produced by SVD will capture most of the variance in the original STA; residual filters will capture noise in the STA. The first pair of spatial and temporal filters captured a majority of the STA variance and resembled the filtering properties of the RF center. The second pair captured far less structure in the STA and appeared to describe the filtering properties of the antagonistic surround. The third pair captured minimal variance and appeared only to represent noise. Thus, for this example cell type, the spatiotemporal RF was well-approximated by a single pair of spatial and temporal RF filters, with some residuals likely resulting from delayed temporal filtering by the RF surround. (B) Singular value spectrum showing the explained variance for the top spatial and temporal filter pairs at scotopic and photopic conditions for all cell types. Noise in the STA causes explained variance to plateau at small but nonzero values. Data from three retinas as described in Figure 3B. This result was consistent at both light levels, despite changes in the RFs across light levels. The approximate space-time separability across all cell types can be viewed simultaneously in two-dimensional space-time plots of the STAs from each type (Figure 3A). Violations of space-time separability will manifest as diagonal structure in these images, but visual inspection suggests that most of the structure is along the horizontal and vertical axes. We quantified space-time separability of these RFs by finding the variance explained about the full RF by a single pair of spatial and temporal filters (rank 1 RF; Figure 3B). The first filter pair explained >85% of the variance for all six cell types at both rod and cone-mediated light levels.

**Figure 4-supplemental figure 1:**
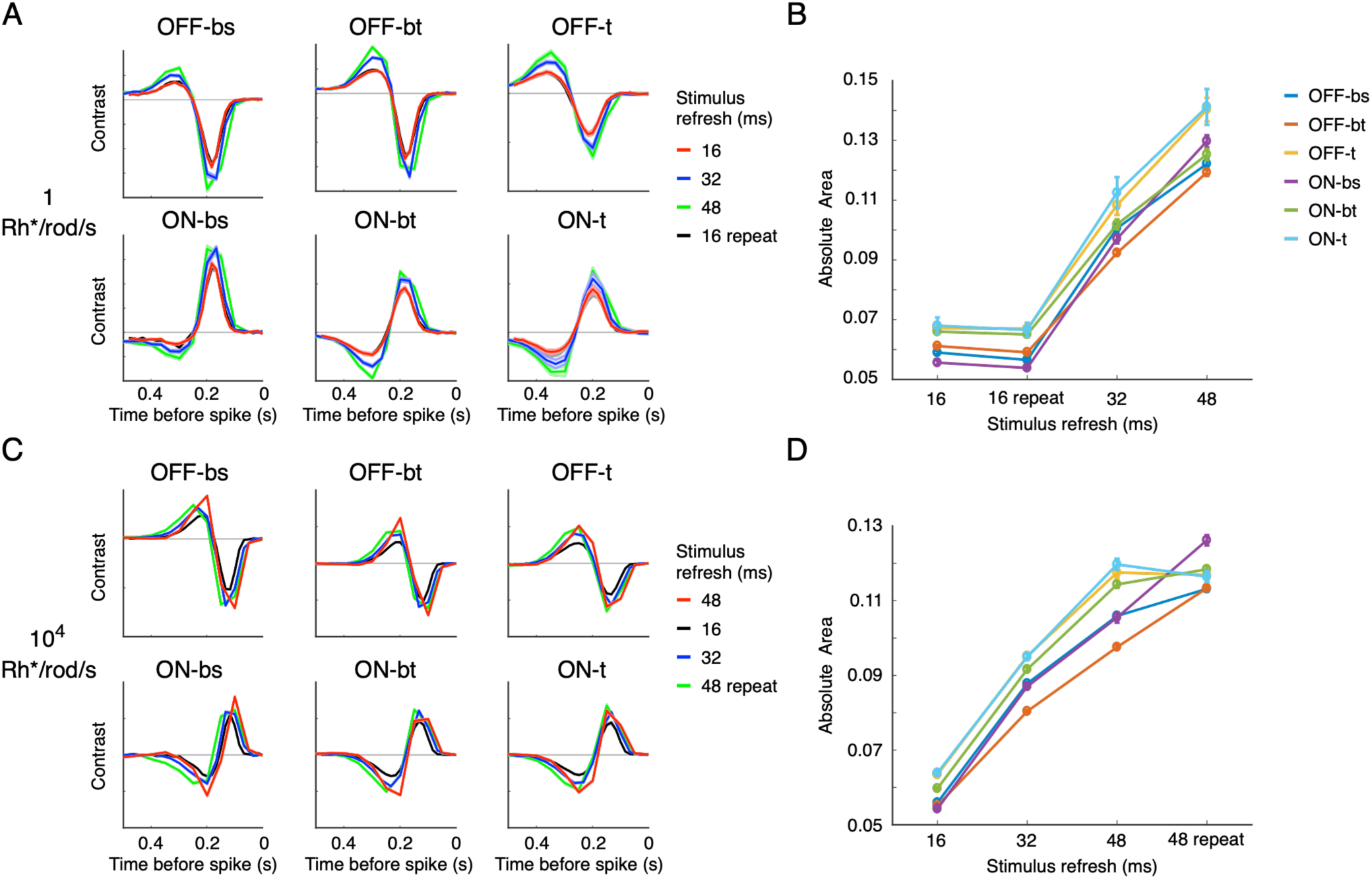
Controls for effective stimulus contrast on temporal RF measurements. Here we control for the impact of contrast adaptation on RF estimation. We have focused on the effects of adaptation to mean stimulus intensity. However, contrast adaptation could also contribute, as altering contrast can change temporal RFs. While each stimulus was presented with the same absolute contrast (variability in the intensity of the checkerboard pixels) across light levels, the effective contrast of the stimulus on the RGCs could change due to change in spatial or temporal integration. To understand how different contrast states affect RF organization, we measured temporal RFs under different white noise refresh rates, thereby changing the effective contrast of the stimulus (A) Temporal RFs estimated with different frame refresh rates for checkerboard noise stimulus at 1 Rh*/rod/s. The dynamics of temporal integration are stable across time resolutions. One stimulus refresh was repeated at the end of the experiment to test for reproducibility. Thickness of average curves includes SD. (B) Gain changes (measured by absolute area under temporal RF curves) are approximately linear with stimulus resolution. Error bars are SEM. (C) The same as A at 10,000 Rh*/rod/s. The few observed kinetic differences in temporal RFs are more consistent with recording rundown than contrast adaptation (compare red and green curves). (D) Quantification of gain changes in C. Error bars are SEM. Data are from one retina. The total number of RGCs varied for different stimulus contrasts and light levels. For OFF-bs, the range of recorded RGCs is 7-9; OFF-bt: 15-17; OFF-t: 6-8; ON-bs: 11-13; ON-bt: 14-17; ON- t: 8-12. In summary, the shape changes of temporal RFs vary linearly with effective stimulus contrast, and the impact on different cell types is relatively stable (B & D, R2=0.91 for 10,000 Rh*/rod/s and 0.93 for 1 Rh*/rod/s). Therefore, we do not believe that contrast adaptation impacts our main conclusions.

## Notes

https://github.com/kmruda/RGC-functional-org

https://github.com/pearsonlab/efficientcoding_adaptation

### Competing Interest Statement

The authors have declared no competing interest.

### Summary of Updates

We have added a new analysis that compares the predictions of efficient coding theory to our measured changes in retinal receptive fields across light levels. This new analysis is presented in figure 10. It show, that previous studies of efficient coding theory made overly simplifying assumptions that led to an incorrect prediction for how receptive field structure should change across changing signal-to-noise conditions, analogous to changes in light level. There is also more discussion of our results and their implications for efficient coding theory.

